# Nuclear envelope disruption triggers hallmarks of aging in lung alveolar macrophages

**DOI:** 10.1101/2022.02.17.480837

**Authors:** Nilushi S. De Silva, Guilherme P.F. Nader, Francesca Nadalin, Kevin de Azevedo, Mickaël Couty, Anvita Bhargava, Cécile Conrad, Mathieu Maurin, Charles Fouillade, Arturo Londono-Vallejo, Rayk Behrendt, Lisa Gallwitz, Paul Saftig, Beatriz Herrero Fernández, José María González-Granado, Guillaume van Niel, Alexandre Boissonnas, Mathieu Piel, Nicolas Manel

## Abstract

Aging is characterized by gradual immune dysfunction and increased risk for many diseases, including respiratory infections. Genomic instability is thought to play a central role in the aging process but the mechanisms that damage nuclear DNA in aging are insufficiently defined. Cells that migrate or reside within confined environments experience forces applied to their nucleus, leading to transient nuclear envelope (NE) ruptures. NE ruptures are associated with DNA damage, and Lamin A/C is required to limit these events. Here, we show that Lamin A/C protects lung alveolar macrophages from NE rupture and hallmarks of aging. Lamin A/C ablation in immune cells results in a selective depletion of lung alveolar macrophages (AM) and a heightened susceptibility to influenza infection. Lamin A/C-deficient AM that persist display constitutive nuclear envelope rupture marks, DNA damage and p53-dependent senescence. In wild-type mice, we found that AM migrate within constricted spaces *in vivo*, at heights that induce NE rupture and DNA damage. AM from aged wild-type mice and from Lamin A/C-deficient mice share an upregulated lysosomal signature with CD63 expression, and we find that CD63 is required to clear damaged DNA in macrophages. We propose that induction of genomic instability by NE disruption represents a mechanism of aging in alveolar macrophages.

## Introduction

Age is a major risk factor for a large number of diseases including respiratory viral infections such as influenza virus and SARS-CoV-2 (Schneider et al., 2021). Chronological aging is characterized by gradual immune dysfunction, which limits protective responses (Nikolich- Žugich, 2018). Studies in genetic models have shown that induction of immune dysfunction can contribute to organismal aging, in part through inflammation (Desdín-Micó et al., 2020; Yousefzadeh et al., 2021). At the cellular level, cells accumulate hallmarks of aging which include genome instability, cellular senescence and altered proteolytic systems (López-Otín et al., 2013). Genomic instability has been proposed to play a central role in driving the aging process (Schumacher et al., 2021). However, the intracellular mechanisms that compromise genome stability during aging in immune cells are ill defined.

Immune cells are highly migratory (Luster et al., 2005) and constantly go through narrow spaces that lead to cellular deformation (Pflicke and Sixt, 2009; Raab et al., 2016). There is an emerging recognition that genome instability can be driven by mechanical forces imparted on the nucleus. Constricted or confined microenvironments *in vivo* exert pressure on the cells that migrate through or reside within these spaces (Garcia-Arcos et al., 2019). These forces can deform the nucleus – the largest and stiffest organelle in the cell (Nader et al., 2021a) – leading to transient nuclear envelope ruptures (Denais et al., 2016; Irianto et al., 2017; Raab et al., 2016). These ruptures cause DNA damage as a result of compromised nuclear compartmentalization (Cho et al., 2019; Denais et al., 2016; Earle et al., 2020; Irianto et al., 2017; Raab et al., 2016; Shah et al., 2021; Xia et al., 2018). The consequences of nuclear envelope rupture-induced DNA damage are largely unknown and demonstrate cell-type specificity. It induces, for instance, senescence in a non-transformed epithelial cell line, while it promotes tumor cell invasion in a breast cancer model (Nader et al., 2021b). Nuclear envelope rupture also exposes genomic DNA to the cytosol, which recruits the interferon-inducing cytosolic DNA sensor cGAS (Denais et al., 2016; Raab et al., 2016). Several mechanisms keep cGAS activation in check following nuclear envelope rupture (Gentili et al., 2019; Guey et al., 2020; Lan et al., 2014), thereby limiting interferon responses that can themselves alter genomic stability (Banerjee et al., 2021; Moiseeva et al., 2006; Morales et al., 2017). Among immune cells, nuclear envelope rupture has been observed in neutrophils and dendritic cells *in vitro* (Raab et al., 2016; Thiam et al., 2020). However, the *in vivo* occurrence and consequences of nuclear envelope rupture in the immune system are not known.

The Lamin meshwork, composed of Lamin A/C, B1 and B2, provides essential mechanical protection of the nucleus (De Vos et al., 2011; Vargas et al., 2012). During migration of cell lines in confined spaces *in vitro*, Lamin A/C is required to limit nuclear envelope rupture, DNA damage and cell death (Denais et al., 2016; Raab et al., 2016). In tissues, Lamin A/C was shown to protect against nuclear envelope rupture in the context of the beating heart and skeletal muscles, that are mechanically strained tissues (Cho et al., 2019; Earle et al., 2020). In humans, mutations in the Lamin A/C gene cause diverse and severe disease manifestations, including accelerated aging phenotypes, and some of these mutations have been associated with decreased genome stability and increased nuclear envelope rupture (De Vos et al., 2011; Earle et al., 2020). In immune cells, Lamin A/C expression is regulated and impacts immune functions in disease contexts (Saez et al., 2020). Whether Lamin A/C provides protection against aging in the immune system is not known.

## Results

### Lamin A/C is required to maintain alveolar macrophages (AM) in the lung

To investigate the physiological consequences of nuclear envelope rupture in immune cells, we sought to identify genetic models that would exacerbate it. The Lamin A/C protein is required to limit nuclear envelope ruptures in constricted spaces (Cho et al., 2019; Earle et al., 2020; Nader et al., 2020). We thus hypothesized that if immune cells undergo NE rupture *in vivo*, Lamin A/C depletion should increase its occurrence. To avoid confounding cell-extrinsic effects of Lamin A/C deficiency in non-immune cells, we analyzed the spleen, lymph nodes, bone marrow and lungs of mice with immune-specific ablation of Lamin A/C (*Lmna*^fl/fl^ Vav1-Cre^+/-^, referred to as Lamin A/C CKO hereafter (de Boer et al., 2003; Kim and Zheng, 2013)). We did not detect a loss of a range of major immune populations including dendritic cell, macrophage, T, B, natural killer cells and hematopoietic stem cell populations in the spleen, lymph nodes and bone marrow (**Figure S1A- C**). However, in the lung we found that alveolar macrophages (AMs) are specifically depleted in Lamin A/C CKO mice (**Figure 1A**), whereas the frequency of lung dendritic cells populations, eosinophils and interstitial macrophages was unperturbed (**Figure S2A, S2B**). This depletion of AM was also confirmed upon myeloid-specific ablation of Lamin A/C (**Figure S2C**).

**Figure 1.**
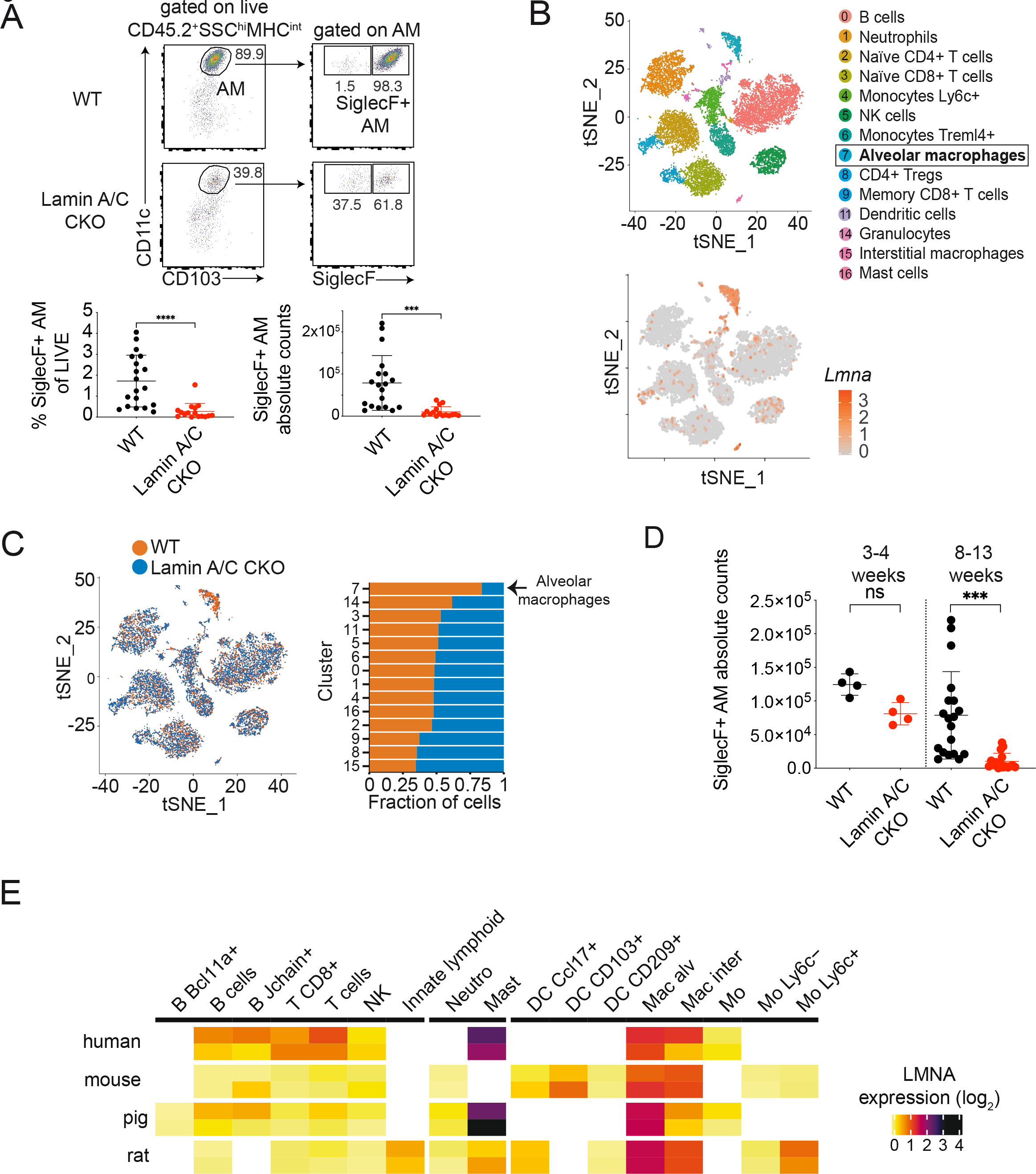
Lamin A/C protects alveolar macrophages from depletion. **(A)** Flow cytometric analysis of AM in WT and Lamin A/C CKO lungs. Top, representative samples. Bottom, percentage of live and absolute counts of AM (n = 16-19 mice combined from 10 independent experiments, bar indicates mean ± SD unpaired t-test). **(B)** tSNE representation of cells identified via single cell RNA sequencing (scRNAseq) of CD45.2+ lung immune cells sorted from WT and Lamin A/C CKO mice (n = 2 female mice per genotype). Top, cluster labelling. Bottom, normalized *Lmna* expression. **(C)** Annotation of WT and Lamin A/C CKO cells in each cluster. Left, tSNE representation. Right, fraction of WT and Lamin A/C CKO per cluster. **(D)** Absolute counts of AM (CD45.2^+^SSC^hi^MHC^int^SiglecF^+^) in lung of WT vs Lamin A/C CKO lung based on flow cytometric analysis at 3-4 and 8-13 weeks of age (n = 4-19 mice, combined from 12 independent experiments, one-way ANOVA with Šidák). **(E)** Expression of the Lamin A/C-coding gene in clusters of lung cells from human, rat, mouse and pig.

To determine if AM were reduced in Lamin A/C CKO as a result of cell loss or phenotypic changes, we performed single cell RNA sequencing (scRNAseq) on CD45.2^+^ cells sorted from the lungs of WT and Lamin A/C CKO mice. Clustering analysis identified immune populations known to populate murine lung (**Figure 1B, top, Figure S3A, S3B, S3C, Table S1**). AM were found to have the highest level of Lamin A/C expression within the immune compartment of the lung, suggesting a particular dependency of this cell type on high Lamin A/C expression (**Figure 1B, bottom**). Indeed, we found that Lamin A/C CKO cells were depleted within the AM cluster in comparison to WT, whereas WT and Lamin A/C CKO cells were equally distributed among the other immune clusters (**Figure 1C**). Lamin A/C CKO mice analyzed at 3-4 weeks of age did not have a strong depletion of AM, in contrast to 8-13 week old mice, suggesting a progressive loss (**Figure 1D**). High levels of Lamin A/C were also observed in AMs of human, pig and rat (**Figure 1E, Table S2**) (Raredon et al., 2019). Our data therefore demonstrate that Lamin A/C is required for the presence of AM, but not other immune cells, within the lung immune compartment.

### A p53-dependent response to DNA damage drives AM depletion

To determine how Lamin A/C deficiency leads to depletion of AM, we examined enriched signatures in differentially expressed genes (DEGs) (**Table S3**). We found an enrichment for p53 targets and for genes involved in apoptotic signaling in response to DNA damage, in genes upregulated in Lamin A/C CKO AMs (**Figure 2A, S4A)**. We also found that Lamin A/C CKO AMs have a higher level of the DNA damage marker γH2AX at steady-state than that observed in WT littermates (**Figure 2B**). In addition, these cells were more sensitive to etoposide, an inducer of DNA damage, applied *ex vivo* (**Figure 2C**).

**Figure 2.**
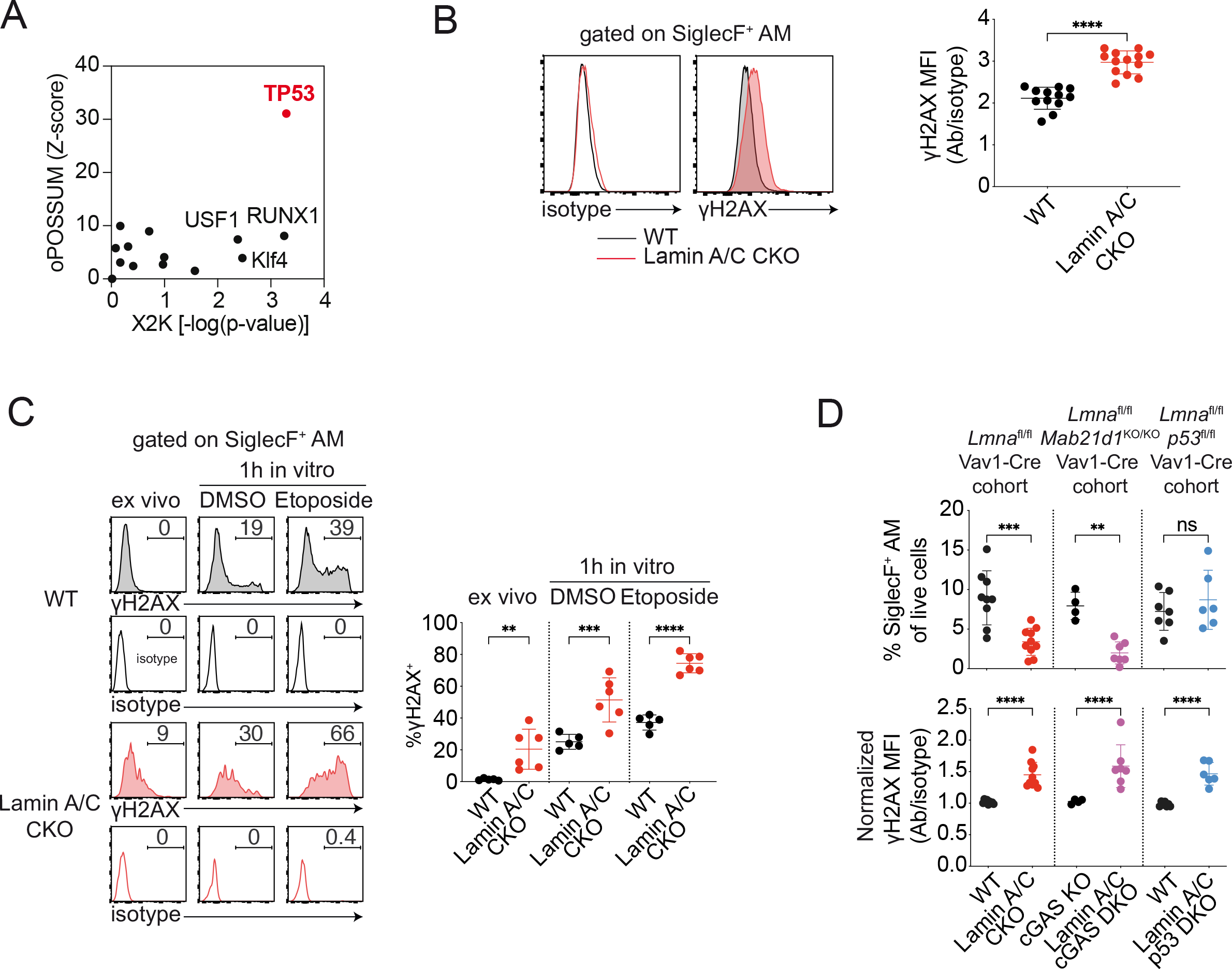
Loss of alveolar macrophages is caused by a p53-dependent response to DNA damage. **(A)** Transcription factor binding site enrichment in promoters of differentially expressed genes (DEGs) upregulated in Lamin A/C CKO and specific to the AM cluster, using oPOSSUM and X2K methods. **(B)** Intracellular levels of γH2AX within WT and Lamin A/C CKO AM (CD45.2^+^SSC^hi^MHC^int^SiglecF^+^). Left, representative samples. Right, mean fluorescence intensity (MFI) of γH2AX signal normalized to isotype control signal (n = 12-13 mice combined from 11 independent experiments, bar indicates mean ± SD, unpaired t test). **(C)** γH2AX response to DNA damage in AM (CD45.2^+^SSC^hi^MHC^int^SiglecF^+^). WT vs Lamin A/C CKO lung single cell suspensions were analyzed directly or incubated with DMSO or 50 μM etoposide for 1 hour. Left, representative samples. Right, quantification of γH2AX^+^ cells (n = 5 or 6 mice, combined from 3 independent experiments, bar indicates mean ± SD, one-way ANOVA with Šidák test). **(D)** Impact of cGAS and p53 deficiencies on AM percentages and γH2AX levels in Lamin A/C CKO mice. 3 breeding cohorts are shown: WT (*Lmna*^fl/fl^ Vav1-Cre^-/-^) vs Lamin A/C CKO (*Lmna*^fl/fl^ Vav1-Cre^+/-^), cGAS KO (*Lmna*^fl/fl^ *Mab21d1*^KO/KO^Vav1-Cre^-/-^) vs Lamin A/C cGAS DKO (*Lmna*^fl/fl^ *Mab21d1*^KO/KO^Vav1-Cre^+/-^) and WT (*Lmna*^fl/fl^ *p53*^fl/fl^Vav1-Cre^-/-^) vs Lamin A/C p53 DKO (*Lmna*^fl/fl^ *p53*^fl/fl^Vav1-Cre^+/-^). In each experiment, cGAS KO and Lamin A/C cGAS DKO or WT and Lamin A/C p53 DKO were analyzed together with age-matched WT and Lamin A/C CKO mice. Top, fraction of AM. Bottom, γH2AX MFI is first normalized to isotype control level in each sample, and next to WT analyzed in the same experiment (n = 4-10 mice combined from 8 independent experiments, bar indicates mean ± SD, one-way ANOVA with Šidák test).

We thus asked if AM depletion was due to senescence associated with DNA damage. DNA damage can induce cellular senescence (Gorgoulis et al., 2019). Expression of senescence-associated cell cycle inhibitor p21 was increased in Lamin A/C CKO AM (**Figure S4B, Table S3**). The dsDNA sensor cGAS can promote senescence by detecting damaged self DNA accumulating in the cytosol (Gluck et al., 2017; Yang et al., 2017). We generated mice with double knockout of Lamin A/C and cGAS (*Lmna*^fl/fl^ *Mab21d1*^KO/KO^Vav1-Cre^+/-^, referred to as Lamin A/C cGAS DKO). Removal of cGAS did not rescue AM depletion in Lamin A/C CKO mice and elevated DNA damage persisted (**Figure 2D**). In agreement with this result, we did not detect expression of cGAS- responsive inflammatory cytokines in Lamin A/C CKO AM such as IL6, IL1β and IL8 (**Table S3**)

(Gluck et al., 2017; Yang et al., 2017). Since p53 induces cellular senescence in response to DNA damage (Reinhardt and Schumacher, 2012), we generated double knockout of Lamin A/C and p53 in immune cells (referred to as Lamin A/C p53 DKO). Conditional knockout of p53 was utilized to delay the onset of tumor growth and create a time window to analyze AM in tumor-free conditions (DeMicco et al., 2013). The depletion of AM was rescued in Lamin A/C p53 DKO mice, despite γH2AX levels remaining elevated in the surviving cells (**Figure 2D, Figure S4C**). These findings demonstrate that AM depletion in Lamin A/C CKO mice is a consequence of the response to DNA damage through p53 and is independent of cGAS.

### Alveolar macrophages (AM) migrate in constricted spaces in the lung

Lamin A/C CKO AMs are progressively lost, consistent with an accelerated senescence process triggered after birth (**Figure 1D**). At steady state, most AM are thought to be sessile and stick to the alveolar epithelium. However slow displacement events within the same alveola or between alveoli through the pores of Kohn have been observed (Neupane et al., 2020). Interestingly, pores of Kohn are infrequent in newborns and increase in number during the first few weeks of life (Amy et al., 1977). Furthermore, we noticed that AM located inside pores of Kohn appeared to have deformed nuclei (Neupane et al., 2020). To determine if AM migrate in the lung at heights that induce nuclear envelope rupture, we followed their movements by live imaging in WT mice. We identified several examples of cell squeezing consistent with entry into pores of Kohn (**Figure 3A and Movie 1**). At the most acute point of squeezing, we measured the confined width of cells to be approximately 2–3 μm (**Figure 3B, Movie 2, Figure S5A**). These observations demonstrate constricted migration of AM *in vivo* within the lung, suggesting a potential relationship between this migration and the onset of the senescence process in Lamin A/C CKO AM.

**Figure 3.**
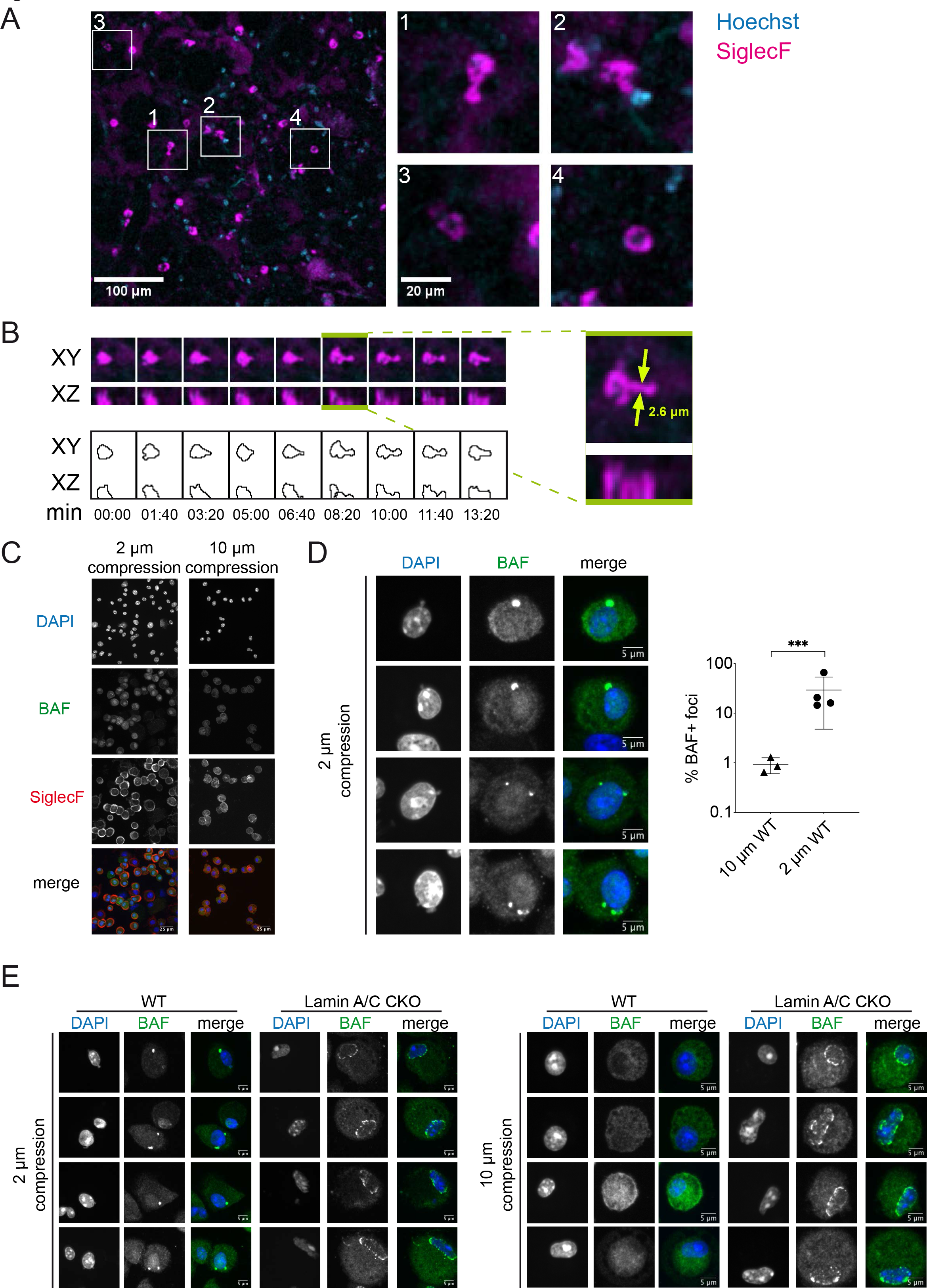
Constricted migration and nuclear envelope rupture in alveolar macrophages. **(A)** Live imaging of WT mouse lung (female, 12 weeks old) after administration of Hoechst and an anti-SiglecF antibody. Left, broad field of a lung region. Right, individual SiglecF^+^ AM demonstrating constricted migration (see also Movie 1). **(B)** Time-lapse of an AM undergoing constricted migration. Top left, images of XY and XZ planes overt time. Bottom left, contour representation. Right, measurement of the cell width at the most acute point of squeezing. **(C)** Cells from the bronchialveolar lavage (BAL) of WT mice confined at a height of 2 µM and 10 μm for 1.5 hours and subsequently stained for DAPI, BAF and SiglecF (representative of n = 4 independent experiments, each time BAL was pooled from 2 mice aged 8-26 weeks). **(D)** BAF foci in AM (SiglecF^+^) from WT BAL following confinement for 1.5 hours. Left, representative images at 2 µM height. Right, quantification of the percentage of BAF foci+ AM after confinement at 2 µm or 10 µM (n = 3-4 independent experiments, each time BAL was pooled from 2 mice aged 8-26 weeks, bar indicates mean ± SD, unpaired t test on log-transformed data). **(E)** BAF staining in AM (SiglecF^+^) from WT and Lamin A/C CKO BAL, following confinement at 2 μm and 10μm for 1.5 hours (representative of n = 3-4 experiments, each time BAL was pooled from 2 mice aged 8-26 weeks). Different scale bars were used between 2 μm and 10μm to accommodate for cell spreading at 2 μm.

### Lamin A/C protects against accumulation of nuclear envelope rupture marks in AM

A wide range of cell types display nuclear envelope rupture when passing through microchannels with 2 μm constrictions or following confinement at heights of 2 μm *in vitro* (Denais et al., 2016; Raab et al., 2016). To determine if this height also induces rupture in AM, we isolated bronchoalveolar lavage (BAL) cells from WT mice and confined these cells at 2 μm and 10 μm, a control height that does not lead to nuclear deformation. SiglecF staining confirmed the identity of AM in cells from the BAL (**Figure 3C**). Upon nuclear envelope rupture, the DNA-binding protein Barrier-to-Autointegration Factor (BAF) forms characteristic foci on DNA herniations protruding from the nucleus (Denais et al., 2016; Halfmann et al., 2019). We identified a fraction of cells at 2μm that displayed nuclear DNA blebbing which was associated with foci of BAF (**Figure 3D**). In contrast, nuclear blebbing and BAF foci were largely absent in AM confined at 10 μm.

Next, we examined nuclear envelope rupture in AM from Lamin A/C CKO mice following BAL cell confinement at 2 μm and 10 μm (**Figure S5B**). We confirmed Lamin A/C ablation in AM obtained from BALs at the protein level (**Figure S5C**). Consistent with a role of Lamin A/C in nuclear shape, Lamin A/C CKO AM confined at 2 μm were characterized by nuclei with reduced circularity and roundness and increased aspect ratio compared to WT (**Figure S5D**). Strikingly, a discontinuous ring of BAF foci was present at the nuclear rim of Lamin A/C CKO AM (**Figure 3E**). This discontinuous ring of BAF foci was present also at 10 μm (**Figure 3E**), suggesting that it was not induced by 2 μm confinement but was present constitutively. In HeLa cells, Lamin A/C depletion is not sufficient to induce such a ring and instead favors BAF diffusion (Haraguchi et al., 2008). Therefore, the discontinuous ring of BAF foci that we observed in Lamin A/C CKO is rather in agreement with the cells undergoing repetitive nuclear envelope rupture and accumulating multiple individual BAF foci. Lamin A/C therefore protects AM from accumulation of nuclear envelope rupture marks.

### DNA damage in AM upregulates CD63 and a lysosomal signature

We next examined the functional consequences of Lamin A/C deficiency and DNA damage in AM. Differential gene expression analysis, together with a bootstrap validation step, was used to identify the most robustly affected DEGs (**Figure 4A**). We identified a downregulation of ribosomal genes that was common among clusters. We also detected upregulation of a set of genes specifically in the remaining AM of Lamin A/C CKO mice. The second most induced gene was *Cd63*, a tetraspanin that is associated with endosomal and lysosomal membranes. We validated increased protein expression of CD63 in Lamin A/C-knockout AM via intracellular flow cytometry (**Figure 4B**). We were also able to validate at the protein level the reduced expression of CD88, also known as complement component 5a receptor 1 or C5AR1 (**Table S3, Figure S6A**). We also identified upregulated expression of a number of proteases including the macrophage elastase (*Mmp12*) and cathepsins (*Ctsk*, *Ctsd*, *Ctsl*, *Ctsz*, *Ctsb*) (**Figure 4A, Table S3**). We confirmed increased Cathepsin L activity in the BAL of Lamin A/C CKO mice (**Figure S6B**). The BAL of Lamin A/C CKO mice also contained elevated levels of neutrophils and T cells (**Figure S6C**) indicative of altered alveolar homeostasis.

**Figure 4.**
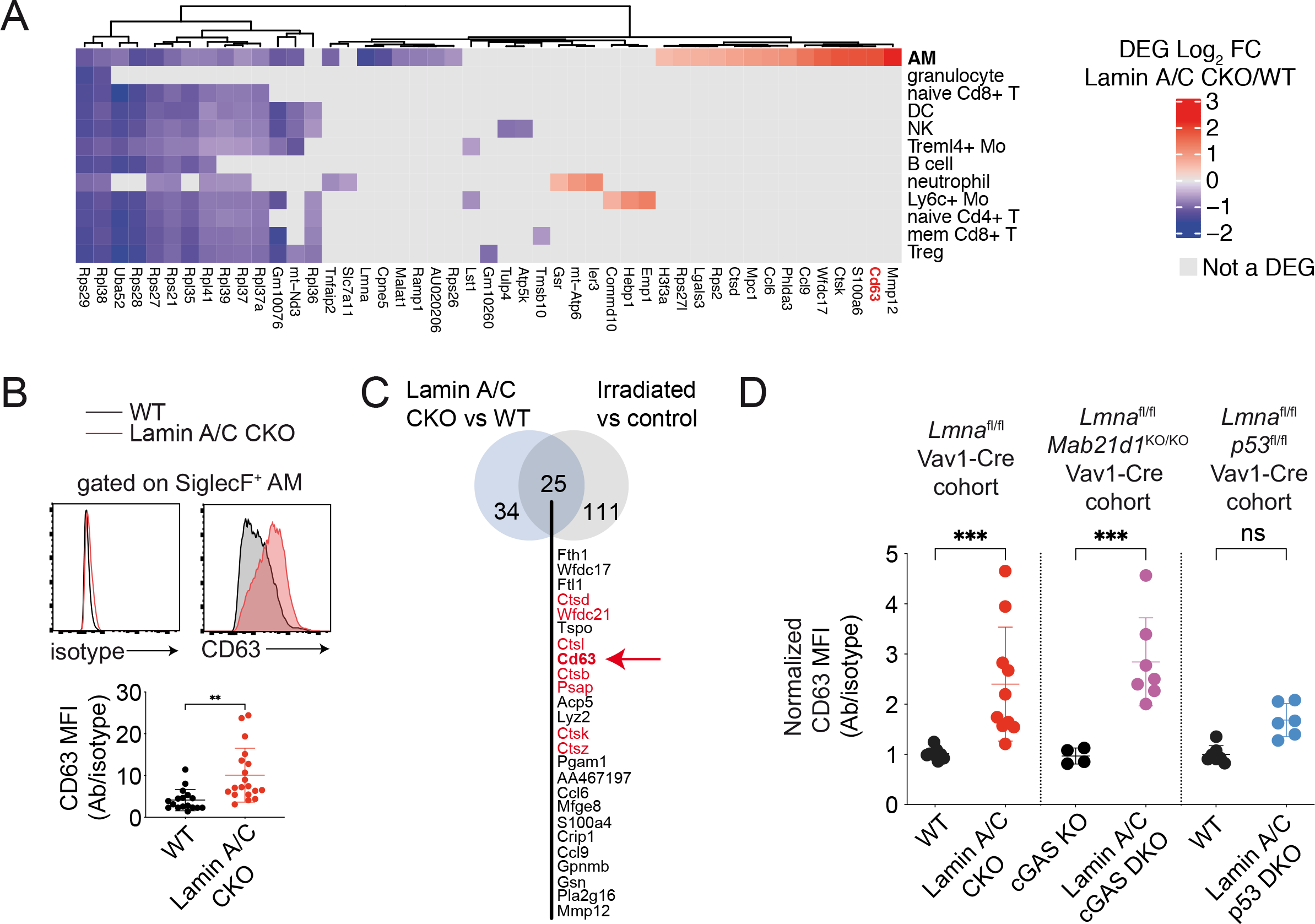
A lysosomal/CD63 response is induced by DNA damage. **(A)** Heatmap showing the significant DEGs in Lamin A/C CKO vs WT cells within cell clusters. Cell clusters are ordered by decreasing fraction of WT cells. Genes were clustered with a complete linkage method computed on the Manhattan distances between genes, shown as dendrogram. **(B)** Intracellular levels of CD63 in lung AM (CD45.2^+^SSC^hi^MHC^int^SiglecF^+^). Top, representative samples. Bottom, CD63 MFI normalized to isotype control (n = 17-19 mice combined from 15 independent experiments, bar indicates average ± SD, unpaired t test). **(C)** Intersection of the DEGs identified in Lamin A/C CKO vs WT AM and the DEGs identified in AM 5 months post-17 Gy irradiation vs no irradiation (control). Cd63, Psap and cathepsin genes are highlighted in red. **(D)** Intracellular levels of CD63 in lung AM (CD45.2^+^SSC^hi^MHC^int^SiglecF^+^) from 3 breeding mouse cohorts: WT (*Lmna*^fl/fl^ Vav1-Cre^-/-^) vs Lamin A/C CKO (*Lmna*^fl/fl^ Vav1-Cre^+/-^), cGAS KO (*Lmna*^fl/fl^ *Mab21d1*^KO/KO^Vav1-Cre^-/-^) vs Lamin A/C cGAS DKO (*Lmna*^fl/fl^ *Mab21d1*^KO/KO^Vav1- Cre^+/-^) and WT (*Lmna*^fl/fl^ *p53*^fl/fl^Vav1-Cre^-/-^) vs Lamin A/C p53 DKO (*Lmna*^fl/fl^ *p53*^fl/fl^Vav1-Cre^+/-^) (n = 4-10 mice combined from 8 independent experiments, bar indicates mean ± SD, one-way ANOVA with Šidák test).

To test if the transcriptional signature of Lamin A/C CKO was the result of an initial DNA damaging event, we analyzed a single-cell RNA seq dataset of whole lung, five months post-irradiation, which is associated with lung fibrosis. We identified upregulated DEGs in the AM cluster of irradiated mice (**Table S4**) and computed the overlap with the upregulated DEGs of Lamin A/C CKO AM. Among the genes that overlapped between these two scRNAseq datasets, were *Cd63*, lysosomal prosaposin (*Psap*), *Wfdc21* and cathepsins (**Figure 4C, Figure S6D**). We next tested if DNA damage itself or the signaling response to DNA damage was inducing CD63, using Lamin A/C p53 DKO AM. In contrast to Lamin A/C CKO, elevation of CD63 expression was blunted in Lamin A/C p53 DKO AM (**Figure 4D**). We therefore conclude that AM upregulate CD63 in Lamin A/C CKO AM in a p53-dependent manner, and in response to irradiation in WT AM. Together with our previous results, this suggests that Lamin A/C deficiency causes DNA damage in AM, leading to p53 activation, which induces CD63 and senescence.

### CD63 is required for damaged DNA clearance in macrophages

CD63 is a tetraspanin that has been proposed to participate to a wide range of intracellular trafficking processes (Pols and Klumperman, 2009). However, the function of CD63 in macrophages is not known. CD63 and lysosomes have been associated with genomic DNA accumulating in the cytosol following the induction of DNA damage in tumor cell lines (Shen et al., 2015) and in senescent regions of human fibrotic lung (Borghesan et al., 2019). CD63 has also been implicated in loading nuclear content into exosomes following micronuclei formation in cancer cell lines (Yokoi et al., 2019). This raised the possibility that CD63 is implicated in the response to damaged DNA in macrophages. We performed dsDNA staining on bone-marrow derived macrophages and observed an accumulation of cytoplasmic dsDNA in CD63 KO cells compared to WT cells (**Figure 5A)**. To investigate whether CD63 may play a role within the context of damaged DNA in AM, we assessed the level of DNA damage accumulating in CD63 KO AM upon *ex* vivo application of etoposide, a DNA damage inducer. We detected a higher percentage of γH2AX+ AM in CD63 KO compared to WT in response to etoposide (**Figure 5B, Figure S6E**). Taken together, these data reveal that CD63 is required to limit the accumulation of DNA damage and clear cytoplasmic DNA in macrophages.

**Figure 5.**
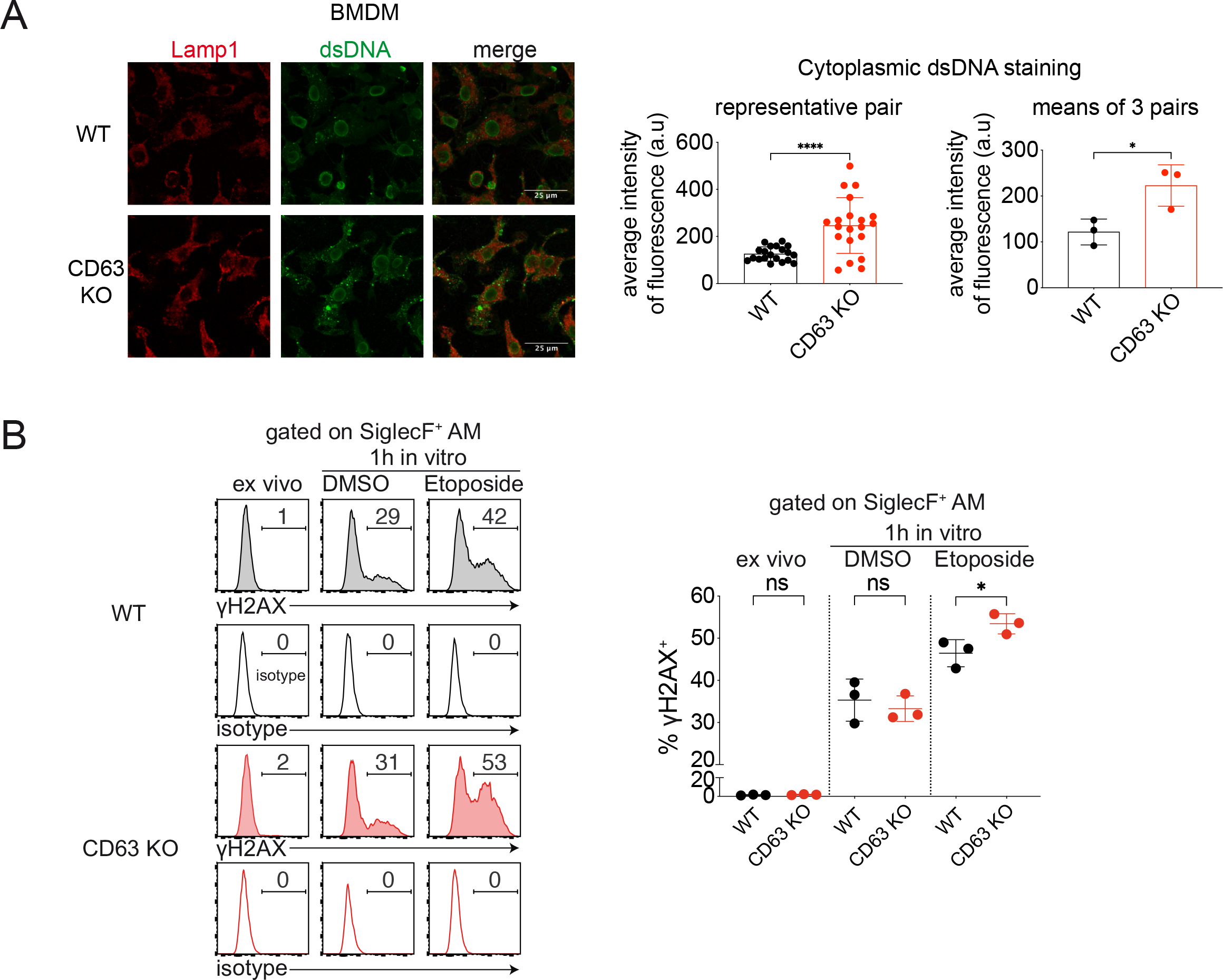
CD63 is required for clearance of damaged DNA in macrophages. **(A)** Lamp1 and dsDNA staining in none marrow-derived macrophages (BMDM) from WT and CD63 KO mice. Left, representative images. Middle, average fluorescence intensity of the dsDNA signal per cell quantified outside the nucleus in a representative pair of WT vs CD63 KO (each symbol represents a single cell, Kolmogorov-Smirnov test). Right, average dsDNA signal per mouse (n = 3 mice per genotype in one experiment, 49 weeks old, bar indicates mean ± SD, unpaired t-test). **(B)** γH2AX levels within AM (CD45.2^+^SSC^hi^MHC^int^SiglecF^+^) of WT vs CD63 KO lung single cell suspension, directly ex vivo or after incubation with DMSO or 50 μM etoposide for 1 hour. Left, representative samples. Right, fraction of γH2AX positive cells (n = 3 mice per genotype in one experiment, 19 weeks old, bar indicates mean ± SD, one-way ANOVA with Šidák test).

### Lamin A/C protects against influenza virus infection

Next, we sought to determine whether the AM alterations that characterize the lungs of Lamin A/C CKO mice have physiophathological consequences. AM play a particularly critical protective role in protection against influenza virus infection (Purnama et al., 2014; Schneider et al., 2014). We therefore infected WT and Lamin A/C CKO mice with the mouse-adapted Influenza A strain PR8. Lamin A/C CKO mice had a lower probability of survival than their WT littermates (**Figure 6A**). Lamin A/C CKO mice also had more dramatic weight loss at critical disease timepoints (**Figure 6B**). Significant weight loss in comparison to the day of virus inoculation was initiated a day earlier than WT littermates (**Figure S7A**). We observed similar results when testing a higher dose of Pr8 (**Figure S7B, 7C, 7D**). We conclude that Lamin A/C is required in immune cells to protect against influenza virus infection.

**Figure 6.**
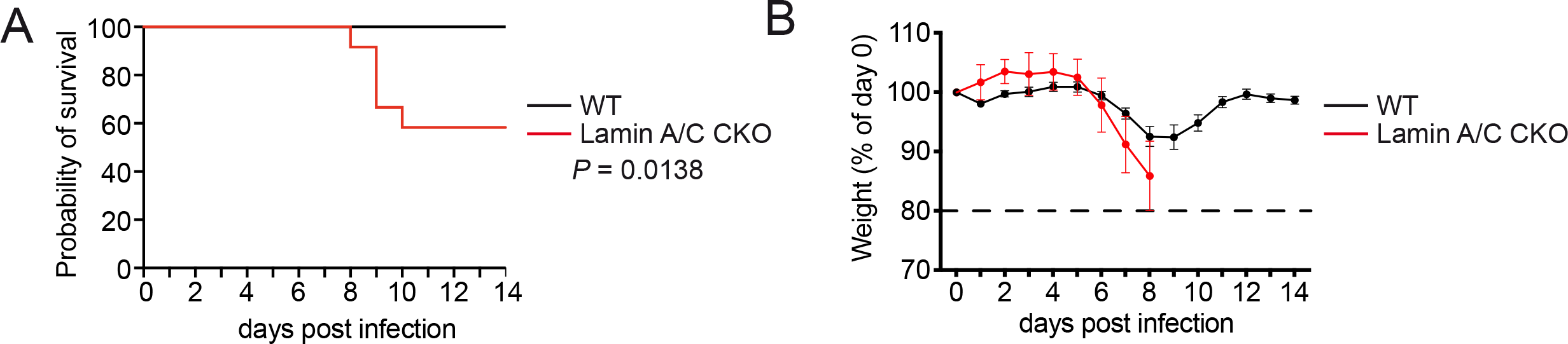
Lamin A/C protects against influenza virus infection. **(A)** Survival of WT vs Lamin A/C CKO mice following infection with 50 pfu of Influenza A The virus PR8 was delivered via the intranasal route (n = 12 female mice per genotype, combined from 2 independent experiments, Log-rank Mantel-Cox test). **(B)** Percentage of day 0 weight observed each day post infection described in (E). Curves of average ± SEM weights are shown and continued until a first death occurs in the group.

### Lamin A/C protects against acquisition of hallmarks of aging in AM

Poor disease outcomes following infection by respiratory viruses, such as influenza virus, in both mice and humans, is a well-characterized consequence of an aged immune system (Schneider et al., 2021). This raised the possibility that Lamin A/C CKO leads to accelerated aging of AM. We showed that WT AM can undergo nuclear envelope rupture at confinement heights that mimic the constricted spaces these cells migrate through *in vivo*. AM are long-lived, self-renewing cells of embryonic origin (Kopf et al., 2015; Sieweke and Allen, 2013). We therefore hypothesized that if repeated rupture events accumulate over time in vivo, WT AM should gradually acquire a signature related to Lamin A/C CKO signature. In mice aged for 63 weeks, we detected a reduction of AM (**Figure S8A**), in agreement with a drop in AM levels previously reported in mice aged for more than 80 weeks (Krishnarajah et al., 2021; Wong et al., 2017; Xin et al., 2015). Next, we analyzed gene expression in AM from mice of increasing age using the Tabula Muris Senis (TMS) scRNAseq atlas, a resource characterizing aging in mouse tissues (The Tabula Muris Consortium et al., 2020). We performed a differentially expression analysis between “young” (1, 3 months) and “old” (18, 21, 30 months) age groups to define a signature of aging in AM (**Figure 7A, Table S5**). Strikingly, *Cd63* was the top upregulated hit in each of the aged timepoints compared to each of the young timepoints. *Cd63* expression was induced between 3 and 18 months of age (**Figure 7B**). We analyzed mice at 14 weeks (young) and 63 weeks (old) of age, that had an identical genetic background and were housed in the same facility. We confirmed at the protein level that aged AM have a higher level of CD63 than their young counterparts (**Figure 7C**). We also conducted a gene set enrichment analysis comparing the TMS signature of aging in AM and the signature of Lamin A/C ablation in AM to appreciate if there was overlap (**Figure 7D, Figure S8B**). The Lamin A/C CKO AM signature was enriched in the AM transcriptome of aged mice and the TMS signature of aging in AM was also reciprocally enriched in the transcriptome of Lamin A/C CKO AM. By intersecting the DEG identified in the 24 months vs 3 months TMS signature of aging in AM with those identified in the Lamin A/C CKO AM signature we observed that along with *Cd63*, *Psap, Wfdc17* and cathepsins were also upregulated in both datasets (**Figure 7E, left**). To confirm this data, we extracted the signature of aging in AM from an independent dataset, the Lung Aging Atlas (LAA) that compared the age points 3 months and 30 months (Angelidis et al., 2019). Using this LAA signature, we also observed that *Cd63, Psap*, *Wfdc17* and cathepsins were again shared between the Lamin A/C AM CKO signature and the LAA aged AM signature (**Figure 7E, middle**). Finally, to appreciate how divergent different signatures of AM aging can be, we compared the TMS signature of aging with the LAA signature of aging in AM (**Figure 7E, right**). Interestingly, the two signatures were not identical, which could be explained by different facilities and genetic backgrounds. However, within the overlap, upregulated expression of *Cd63, Psap*, *Wfdc17* and cathepsins persisted, suggesting that these changes withstand possible background and environmental variables. We did not detect elevated DNA damage at steady state in aged AM at 63 weeks based on γH2AX staining, but susceptibility to DNA damage, that characterizes Lamin A/C CKO AM, was slightly increased in old macrophages (**Figure S8C**). We therefore found that *Cd63,* cathepsin expression and enhanced susceptibility to DNA damage are common features of aging and Lamin A/C ablation in AM.

**Figure 7.**
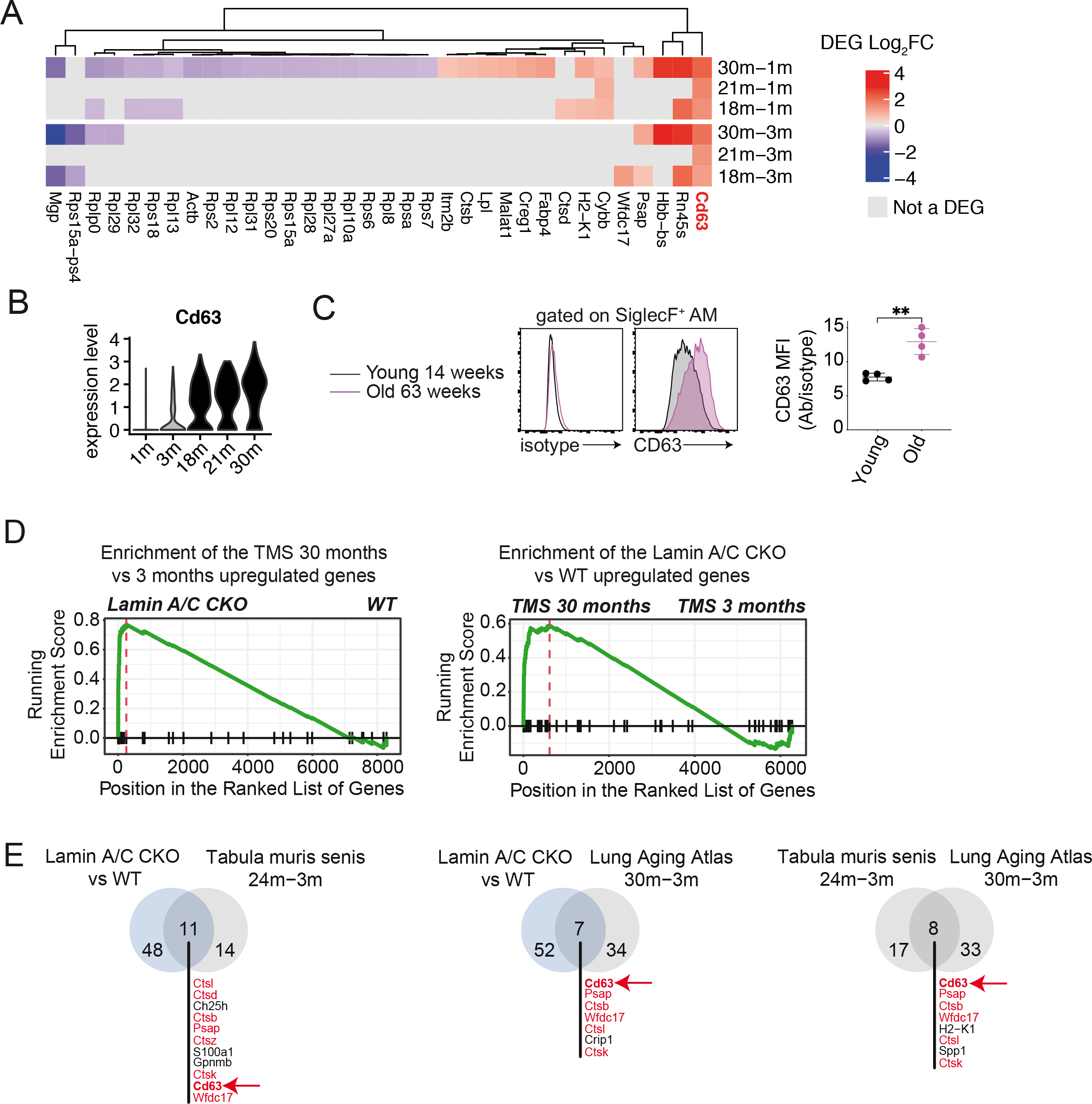
Lamin A/C protects against hallmarks of aging in alveolar macrophages. **(A)** Heatmap showing the significant DEGs in old (18, 21, 30 months) vs young (1 month, 3 months) comparisons in AM in the TMS dataset. Genes were clustered with a complete linkage method computed on the Manhattan distances between genes, shown as dendrogram. **(B)** Violin plot showing the normalized expression of *Cd63* in the AM cluster at different ages. **(C)** Intracellular CD63 levels in AM (CD45.2^+^SSC^hi^MHC^int^SiglecF^+^) of young (14 weeks) vs old (63 weeks) mice. Left, representative samples. Right, CD63 MFI normalized to isotype control (n = 4 mice combined form 2 independent experiments, bar indicates mean ± SD, unpaired t test). **(D)** Reciprocal gene set enrichment analyses, as indicated on figure. **(E)** Intersection of the upregulated DEGs in Lamin A/C CKO vs WT, TMS 24 months vs 3 months and LAA 30 months vs 3 months. Common genes are shown in red and Cd63 is highlighted.

## Discussion

Our results show that AM require Lamin A/C for protection against accumulation of nuclear envelope rupture marks, genome instability and accelerated aging.

Within the lung immune compartment, AM expressed the highest level of Lamin A/C. AM are long-lived cells that self-renew in tissues (Sieweke and Allen, 2013). The longevity of this population *in situ* may necessitate a higher level of nuclear protection to limit the accumulation of DNA damage over time. Lamin A/C levels are thought to scale with tissue stiffness (Swift et al., 2013). Therefore, high Lamin A/C expression in AM may be an adaptation to counter repetitive nuclear deformation and subsequent envelope ruptures caused by constricted migration, such as through pores of Kohn. Such migration may be further stimulated by lung infection (Neupane et al., 2020). Additional mechanical effects, such as constant inflation and deflation that occurs during breathing, could also be a source of biomechanical stress leading to increased nuclear envelope rupture.

Multiple mechanisms likely contribute to the induction of DNA damage during nuclear envelope rupture in AM. In addition to limiting nuclear envelope rupture, Lamin A/C also participates in DNA repair (Gruenbaum and Foisner, 2015). The transient loss of nuclear lamina integrity during rupture in WT cells may thus transiently compromise the repair process. Cytosolic leakage of DNA repair proteins after nuclear envelope rupture may also contributes to DNA damage (Irianto et al., 2017; Xia et al., 2018). Finally, in epithelial and cancer cell lines, the ER-associated exonuclease TREX1 is a driver of this damage after translocation into the nucleus post-rupture (Maciejowski et al., 2015; Nader et al., 2021b). DNA damage caused by nuclear envelope rupture may therefore be the result of a combination of these and other, yet to be determined, molecular events.

Our results provide *in vivo* evidence for a CD63/lysosomal pathway that clears damaged DNA (Shen et al., 2015; Yokoi et al., 2019) and implicate it in the process of aging. CD63 KO AM were more susceptible to DNA damage than WT counterparts, and CD63 KO BMDM constitutively accumulated DNA damage. This indicates that CD63 promotes clearance of damaged DNA in macrophages. Our results also show that CD63 marks macrophages that age over time or through acceleration with increased DNA damage. CD63 induction in Lamin A/C CKO required p53, suggesting that it is a response to genome instability. Accordingly, we found that the CD63/lysosome signature is also induced by exposing lung to ionizing radiation, which is known to induce senescence and accelerate aging, further suggesting that the upstream cause is the presence of DNA damage. While we could show that aged AM are more sensitive to etoposide than young AM, we did not detect increased DNA damage at baseline in aged vs young AM using yH2AX quantification. This suggests that very low yet chronic DNA damage, below the threshold of yH2AX induction or detection, accumulates over-time to induce a CD63/lysosomal response. In support of this idea, whole-lung analyses have detected an increase in DNA damage with age (Braidy et al., 2011; Lee et al., 2021) and an accumulation of DNA damage with age is emerging as a generalized feature of aging (Schumacher et al., 2021).

Our results indicate that disruption of Lamin A/C results in endogenous DNA damage that is sufficient to induce the CD63/lysosomal signature of aged AM. Cell-extrinsic mechanisms likely play an additional role in AM aging. The lung microenvironment has been implicated in age-related changes occurring in AM (McQuattie-Pimentel et al., 2021) and lung tissue mechanics evolve during aging (Sicard et al., 2018). Phagocytosis of DNA or other debris accumulating in the lung spaces could also contribute to lysosome congestion, reducing the ability of AM to clear their own damaged DNA. Exogenous DNA damaging events can also contribute to inducing the AM aging signature, as we found with lethal irradiation. We speculate that additional exogenous events, such as inhaled agents or infection, could also serve as source of exogenous DNA damage. Finally, we also detected increased Cathepsin L activity and immune cell infiltrate detected in the airspaces of mice with immune-specific ablation of Lamin A/C, which could be triggered initially by an AM-intrinsic defect, but subsequently exacerbated by cell-extrinsic responses to a changing microenvironment.

While the progressive acquisition of senescent AM during aging is detrimental, as shown by higher susceptibility to influenza infection in Lamin A/C CKO mice, senescence induction in AM may be beneficial early in life, in agreement with the antagonistic pleiotropy hypothesis (Blagosklonny et al., 2009). For example, it might be a safety mechanism to prevent expansion of damaged cells that self-renew.

Advanced age is a major risk factor for the development of a range of lung diseases. These include conditions such as chronic obstructive pulmonary disease, lung cancer and respiratory infections such as influenza and most pressingly, SARS-CoV-2 (Schneider et al., 2021). AMs play central roles in these diseases (Casanova-Acebes et al., 2021; Purnama et al., 2014; Schneider et al., 2014). We propose that Senescence INduction by Nuclear Envelope Rupture (Nader et al., 2021b) (SINNER) is a mechanism of aging in lung alveolar macrophages. Targeting nuclear envelope rupture may thus constitute a novel opportunity for therapeutic intervention in disease and aging. Beyond AM, we propose that SINNER may be a mechanism of aging in other cell types, that remain to be uncovered.

## Acknowledgements

We thank Helena Izquierdo Fernández and Gisèle Bonne for discussions, Olivier Lantz, Aurélie Darbois Delahousse, Julie Helft and Leila Perie for technical help, Tabula Muris Senis consortium for early access to data and the Institut Curie Institut Curie flow cytometry, NGS and animal facilities. This work was supported by <u>Institut Curie, INSERM, CNRS, Agence Nationale de la Recherche grant LUSTRA (AL), Agence Nationale de la Recherche grants ANR-17-CE15-0025-01, ANR-19-CE15-0018-01, ANR-18-CE92-0022-01, 11-LABX-0043 (NM),</u> Institut National du Cancer grant 2019-1-PL BIO-07-ICR-1 (NM and MP), Institut Thématique Multi-Organisme Cancer grant 19CS007-00 (NM and MP), Fondation chercher et Trouver (NM), Mutuelle Bleue (NM), Fondation ARC pour la recherche sur le cancer grant PGA1 RF20190208474 (GvN), Fondation pour la Recherche Médicale grant EQU202103012774 (NM), EDF grants ORG20190038 and CT9818 (AL), Instituto de Salud Carlos III grant PI17/01395, PI20/00306 (JMGG), imas12 grant i12-AY201228-1 (JMGG), Fondo Europeo de Desarrollo Regional grant “A way to build Europe” (JMGG), EMBO ALTF grant 1298-2016 (NDS), European Commission Horizon 2020 grant H2020-MSCA-IF-2016 DCBIO (NDS), Région Ile-de-France DIM Longévité et Vieillissement (NDS), MCNU FPU program grant FPU18/00895 (BHF).

## STAR Methods

### Mice

All animals were used according to protocols approved by Animal Committee of Curie Institute CEEA-IC #118 and maintained in pathogen-free conditions in a barrier facility. Experimental procedures were approved by the Ministère de l’enseignement supérieur, de la recherche et de l’innovation (APAFIS#24911-2020033119476092-v1) in compliance with international guidelines. C57BL/6JMb21d1tm1d(EUCOMM)Hmgu (Mb21d1^KO/KO^) mice were obtained from The Jackson Laboratory. *Lmna*^fl/fl^ Vav1-Cre^+/-^ (Lamin A/C CKO mice, (de Boer et al., 2003; Kim and Zheng, 2013)) and Lmna^fl/fl^ LysM-Cre^+/-^ (Clausen et al., 1999) were obtained from José María González Granado. p53^flox/flox^ mice (Jonkers et al., 2001) were obtained from Renata Basto. CD63^KO/KO^ were obtained from Paul Saftig (Schröder et al., 2009). The sex of mice used in the experiments shown are listed in **Table S6**. Littermate controls were used in each experiment. The age range was 6-13 weeks unless otherwise stated. Female C57BL/6J elderly mice and young controls were purchased from Charles River Laboratories. Unless otherwise stated, WT controls are *Lmna*^fl/fl^ Vav1-Cre^-/-^. For analysis of DKO animals (Lamin A/C cGAS DKO, Lamin A/C p53 DKO), Lamin A/C CKO of the same age were analyzed with DKO animals in each experiment, together with controls from each cohort.

### Lung preparation for FACS analysis

Dissected lungs were mechanically disrupted using scissors and digested for 30 minutes with 0.4mg/mL collagenase (Sigma C2139) + 20μg/mL DNAse I (Sigma 10104159001) in RPMI at 37 degrees in 12 well tissue culture plates. This was followed by homogenization of the tissue mix over a 100μm filter using 1% BSA/1mM EDTA/PBS and a 5mL syringe plunger. Red blood cells were lysed for five minutes at 4 degrees using red blood cell lysis buffer (Ozyme 42031) and the cells were washed and filtered again over a 100μm filter. The resulting single cell lung suspension was then used in FACS analysis (see **Table S7** for a list of antibodies used in the study) and cell sorting.

### Spleen and lymph node preparation for FACS analysis

Dissected spleen was injected with 0.125mg/mL Liberase (Sigma 5401020001) + 20μg/mL DNAse I (Sigma 10104159001) and inguinal and brachial lymph nodes were disrupted using scissors and incubated with the same Liberase/DNAseI mix. The organs were digested for 20 minutes with Liberase/DNAseI at 37 degrees in 12 well tissue culture plates. This was followed by homogenization of the tissue mixes over a 40μm filter using 1% BSA/1mM EDTA/PBS and a 5mL syringe plunger. Red blood cell lysis was performed as described above and the cells were washed and filtered again over a 40μm filter. The resulting single cell suspensions were used in FACS analysis.

### Bone marrow extraction

Tibias and femurs were cleaned and cut at the ends. Bone marrow was flushed via centrifugation at 2000 rcf for 20 seconds at 4 degrees. Extracted bone marrow was then used directly to generate BMDMs or red blood cell lysis was performed as described above and the cells were washed and filtered over a 40μm filter. The resulting single cell suspension was used in FACS analysis.

### Flow cytometry (FACs) analysis

BAL cells, lung, spleen, lymph node or bone marrow single cell suspension was first stained using a Fixable Viability Dye (ebioscience 65-0865-14) for 20 minutes at 4 degrees in PBS. The cells were then washed with 1% BSA/1mM EDTA/PBS (FACs buffer) and stained with antibodies specific for surface antigens for 30 minutes at 4 degrees in FACs buffer. If intracellular staining was also performed, the Foxp3 Transcription Factor Staining Buffer Set (eBioscience 00-5523-00) was used following the manufacturer’s instructions. For surface staining of CD88, cells were incubated with Fc block in FACs buffer for 15 minutes at 4 degrees after live/dead staining prior to staining for surface antigens. See **Table S7** for details on antibodies used for cell surface and intracellular staining. Cells were filtered over 40μm filter-cap FACs tubes in FACs buffer just prior to analysis on an LSRII flow cytometer.

### Cell sorting and scRNAsequencing

Lung single cell suspension was stained for FACs analysis as described above (live/dead and surface staining) and CD45.2+ lung immune cells were sorted using a BD FACs Aria III from 2 Lmna^fl/fl^Vav1-Cre^+/-^ and 2 Lmna^fl/fl^ Vav1-Cre^-/-^ littermate controls (all female and 10 weeks old). A 100μm nozzle and 20 psi pressure was used during sorting. Cells were sorted into 20% FBS in RPMI with penicillin-streptomycin, 50μM 2-Mercaptoethanol, 1X non-essential amino acids, 10mM HEPES and 1mM sodium pyruvate added and kept at 4 degrees. After cell counting using a LUNA automated cell counter (Logos biosystems), 9600 CD45.2+ lung immune cells per sample were processed using the Chromium Single Cell 3’ Reagents Kits v3 following the manufacturer’s instructions and sequenced using 25,000 reads per cell on a Ilumina NovaSeq 6000.

### *Ex vivo* etoposide treatment

Lung single cell suspension was divided into three parts. One part was kept at 4 degrees (unstimulated), and the remaining 2 parts were stimulated with DMSO or 50μM etoposide (Sigma E1383) for 1 hr at 37 degrees 5% CO2 in 6 well tissue culture plates. After 1 hour the DMSO and etoposide-treated cells were re-harvested via flushing and these samples together with the unstimulated sample, were stained for FACs analysis as described above (live/dead, surface and intracellular staining).

### Bronchialveolar lavage (BAL)

Mice were euthanized via gentle cervical dislocation under isoflurane-mediated anesthesia. An incision was made at the voice box and a 1mL syringe with a 18G blunt safety transfer needle (Dutscher 303129) attachment was inserted into the trachea. Five, 1mL washes of the lung airspaces were performed using warm 0.5% BSA/2mM EDTA/PBS. For flow cytometry analysis or cell confinement, red blood cells were lysed as described above and the BAL cells were washed and filtered over a 40μm filter. For the measurement of cathepsin L activity in BAL, the first 1mL BAL wash was stored separately and the supernatant was used immediately or aliquoted and frozen for future analysis.

### Multi-photon imaging

Two-photon intravital lung imaging was performed according to Dr Simon Cleary recommendations (Cleary et al., 2020). Following anesthesia with ketamine/xylazine (80/15 mg/kg i.p.), slow intratracheal administration of 50µl of a PBS solution containing (1mg/ml) Hoechst and 1µg of anti-siglecF-PE antibody was performed. Subsequently mice were ventilated with room air plus 1.5% isoflurane at 125 breaths/min at 10 μL/g body weight breath volume (Minivent, Harvard Apparatus), with 2–3 cmH2O positive end-expiratory pressure. A custom-made thoracic window was then inserted into an intercostal incision and the left lung was immobilized against the window with 10-30 mmHg negative pressure. Local temperature was monitored and maintained at 33°C using an incubation chamber. The two-photon laser-scanning microscopy (TPL SM) set-up used was a 7MP (Carl Zeiss) coupled to a Ti: Sapphire Crystal multiphoton laser (ChameleonU, Coherent), which provides 140-fs pulses of near-infrared light, selectively tunable between 680 and 1050 nm and an optical parametric oscillator (OPO-MPX, Coherent) selectively tunable between 1,050 and 1,600 nm. The NLO and the OPO beams were spatially aligned and temporally synchronized using a delay line (Coherent). The excitation wavelength was 820 nm for the NLO beam and 1070 nm for the OPO beam. The system included a set of external nondescanned detectors in reflection with a combination of a LP-600-nm dichroic mirror (DM) followed by LP- 462-nm DM with 417-/60-nm emission filter (EF), LP-500-nm DM with 480-/40-nm EF, LP- 550nm DM with 525-/50-nm and 575/50 nm EFs. Real time movies were performed by imaging every 10s by 5 consecutive 3μm z spacing image stack (total 12μm thickness). For all images the objective was a water immersion, plan apochromat ×20 (numerical aperture = 1).

### Cell confinement

Confinement of BAL cells was performed as described (Nader et al., 2021b). Briefly, a 6-well plate cell confiner was used to confine BAL cells at 2μm and 10μm using coverslips containing a microfabricated layer of PDMS micropillars of defined heights, with the height of the pillars determining the height of spatial confinement of cells placed between the coverslip and 6-well plate surface. The coverslips were applied to the cells using large PDMS pillars attached to a modified 6-well plate cover-lid. These large PDMS pillars served to push the confining coverslips onto the cells to confine them at desired heights on the 6 well glass/plastic bottom plates. Imaging of the cells was performed using the same plates (see below).

### Immunofluorescence of confined BAL

For nuclear shape analysis, BAL cells were stained with Hoescht 33342 (Invitrogen R37605) prior to cell confinement. In 1 of the 3 experiments, the cells were also stained for surface expression of CD11c and SigelcF prior to confinement. The cells were imaged immediately while still confined using an epifluorescence DMI-8 Leica inverted microscope equipped with a Hamamatsu OrcaFlash Camera. For immunofluorescence staining, BAL cells were confined for 1.5 hours at at 37 degrees 5% CO2. Staining was performed in the 6 well plate used during confinement after removal of the pillars/coverslips. The cells were fixed with 4% paraformaldehyde for 20 minutes at room temperature and quenched with 0.1M glycine (Life Technologies) for 10 minutes at room temperature. Permeabilization was performed using 0.2% BSA/0.05% saponin in PBS (IF buffer) for 30 minutes at room temperature. The samples were incubated with primary antibodies in IF buffer + 10% goat serum (Sigma G9023) overnight at 4 degrees in a humidity chamber. After 4 washes with IF buffer the secondary antibodies were incubated for 1 hour at room temperature protected from light. The samples were then washed 5 times with IF buffer and a further 2 times each with PBS and then mounting in Fluoromount-G Mounting Medium with DAPI (Fisher scientific 15596276). Samples were kept at 4 degrees, shielded for light prior to imaging. Imaging was performed using spinning-disc confocal microscope with a Yokogawa CSU-X1 spinning-disc head on a DMI-8 Leica inverted microscope equipped with a Hamamatsu OrcaFlash 4.0 Camera, a Nano-ScanZ piezo focusing stage (Prior Scientific) and a motorized scanning stage (Marzhauser). Both microscopes were controlled by MetaMorph software (Molecular Devices).

A macro was developed to measure nuclear shape of confined BAL cells using ImageJ in which individual nuclear regions were identified after thresholding using the DAPI channel and the Analyze Particles function was run. Three repeats of this experiment were performed, and the third included manual selection of CD11c+SiglecF+ cells. Another macro was developed to quantify the number of γH2AX foci and their intensity in confined BAL cells. Here, individual nuclear regions were identified after thresholding using a Z projection of the DAPI channel and SiglecF- cells were excluded. The number of individual γH2AX foci within nuclei was quantified using the Find Maxima function on a Z projection of the γH2AX channel, and the mean intensity of these spots per nuclei was measured.

### Measurement of Cathepsin L activity

Fresh or thawed BAL supernatant was analyzed using a Cathepsin L Activity Assay Kit (Abcam ab65306) kit following the manufacturer’s instructions and the sample fluorescence was measured using a CLARIOstar microplate reader (BMG LABTECH). Readings were normalized by the volume of BAL retrieved.

### BMDM differentiation and immunofluorescence

Extracted bone marrow cells (see methods above) were plated at 1million/mL on untreated plates in RPMI with 10% FBS, penicillin-streptomycin, 50μM 2-Mercaptoethanol, 1X non-essential amino acids, 10mM HEPES and 1mM sodium pyruvate + 10ng/mL human M-CSF (Miltenyi Biotec 130-096-492). Cells were left undisturbed for 6 days and then harvested following detachment using trypsin. BMDM were first fixed in in 2% PFA for 15 minutes in cell culture medium and then 2% PFA in PBS 1X. The cells were washed 3 times in PBS and quenched with 50mM Glycine in PBS for 20 minutes at room temperature, followed by 3 PBS washes. After quenching, the cells were permeabilized in 0.2% BSA, 0.1% saponin in PBS for 15 minutes at room temperature and blocked in 1% BSA, 0.2% saponin in PBS was 10 minutes at room temperature. Primary and secondary antibodies were incubated in 1% BSA in PBS for 30 minutes shielded from light. Cells was mounted using Immuno-mount (Thermo Scientific). BMDM were imaged on SP5 Confocal microscope (Leica). Quantification of average intensity for cytoplasmic dsDNA staining was performed using ImageJ software by selecting region of interest excluding nucleus from each of the cells imaged for quantification.

### Whole mouse irradiation

Ten to fourteen-weeks old females C57BL/6J mice were purchased from Charles River Laboratories and exposed to whole thorax irradiation at a dose of 17 Gy under 2.5% isoflurane anaesthesia. The irradiation procedure was approved by the ethics committee of the Institut Curie CEEA-IC #118 (Authorization number APAFIS#5479-201605271 0291841 given by National Authority) in compliance with the international guidelines. Mice, kept for five months after irradiation, were weekly monitored and sacrificed if predefined ethical endpoints (i.e. loss of 20% body weight, severe dyspnea) were reached.

### Influenza infection

Influenza A virus PR8 (strain A/Puerto Rico/8/1934 H1N1) was a gift from Olivier Lantz. Intranasal infection with 50 or 100 pfu was performed under Ketamine/Xylazine-mediated anesthesia inside a PSMII hood. Mice were weighed daily and all husbandry was performed underneath a PSMII hood with strict containment measures followed. An ethical endpoint of 20% d0 weight loss was observed in all experiments. Animal care and use for these infection experiments was performed in accordance with the recommendations of the European Community (2010/63 / UE) for the care and use of laboratory animals. Experimental procedures were specifically approved by the ethics committee of the Institut Curie CEEA-IC # 118 (Authorization APAFIS # 32125-2021062516083243 v1 given by National Authority) in compliance with the international guidelines.

### Data analysis

Analysis of FACs data was performed using FlowJo v10.6.2. Statistical analyses were performed using PRISM v9.1.2, the type of test performed is detailed in the relevant figure legends. ImageJ Java 1.8.0_66 was used for analysis of immunofluorescence and intravital data. Figures were compiled using Adobe illustrator v15.1.0

### Bioinformatics analysis

#### Quality check, read alignment, computation of the UMI counts, and cell calling

FASTQ files were obtained from BCF files using cellranger mkfastq command from CellRanger v3.0.2 (Zheng et al., 2017). Sequencing quality was assessed using FastQC v11.8 (https://www.bioinformatics.babraham.ac.uk). cellranger count command (--expect cells 6000) was used to map the reads to the annotated mm10 genome (accession: GCA_000001635.6, gene build: 2016-01), compute UMI counts, and call cellular barcodes.

DoubletDetection (https://doubletdetection.readthedocs.io) was run on the gene-cell expression matrix 50 times and a barcode was labelled as *doublet* (*i.e.*, a droplet containing more than one cell) if it was detected as such (p-value ≤ 10^7^) in at least 40 iterations.

#### Processing of the gene-cell expression matrix

The gene-cell expression matrix for each sample was imported in R using Seurat v3.0.0 (Satija et al., 2015) and normalized as follows: the UMI count of each gene *i* in cell *j* was augmented by 1, divided by the total UMI count for cell *j*, multiplied by 10000, and log-transformed. Doublets and cells with < 200 detected genes (UMI count ≥ 1) were filtered out. Only the genes detected in ≥ 2 cells in at least one sample were retained. The normalized matrices for the four samples were concatenated to obtain a joint normalized matrix *M* and gene expression is standardized (Z-score) to obtain a scaled matrix *M’*.

### Generation of the clustering solution

Feature selection methods and clustering parameters were let vary in order to optimize the definition of the cell populations.

Feature selection was performed on *M* (defined above) using either mean.var.plot or vst method from Seurat. They both model the expression dispersion of each gene in order to detect the candidates that showed the highest variability across cells, called highly variable genes. For mean.var.plot, the average expression was set in [0.1, 0.5] and the scaled dispersion was required to be ≥ *d*, with *d* = 0.5, 1, or 1.5. For vst, the number of top variable genes retained was set to 500, 1000, or 2000, respectively.

A principal component analysis (PCA) was possibly performed on *M’* (defined above) after feature selection. The first 50 principal components (PCs) were calculated using an SVD approximation. The number of top PCs to be retained was automatically defined using the following approach. First, a p-value *ph* was computed for each component *PCh* using the JackStraw method (Seurat, default parameters). To identify the contribution to the standard deviation that was likely due to noise, we computed the standard deviation *σnoise* explained on average by *PC40*,…,*PC50*. The optimal number of top PCs was set to the maximal *h* such that the two conditions *σh* ≥ 1.25*σnoise* and *ph* ≤ 10^−5^ simultaneously hold.

Clustering was done using the Waltman and van Eck algorithm, either on the selected features or on the top *h* PCs, by varying the number of neighbors *k* in {30, 40, 50} and the resolution parameter *r* from 0.1 to 1 in steps of 0.1. Overall, 360 clustering solutions were generated.

### Selection of the optimal clustering solution and identification of cell populations

Given a clustering solution *C*, a silhouette width *silh(j)* was computed for each cell *j* using the *n×n* matrix *D*, where *n* is the number of cells and *Djk* is the euclidean distance between cell *j* and cell *k*, computed on the same space where *C* is calculated (features or PCs). The average silhouette width across a cluster *c* will be referred to as *silh(c)*.

For each solution, the cluster with highest average expression ranking across the alveolar macrophage (AM) markers (Marco, Mrc1, Siglecf, Lmna, Itgax) was defined as the putative AM cluster and labelled as *cAM*. For each feature selection method, with or without PCA, we selected the solution such that *silh(cAM)*+*silh(C)* is maximal (12 solutions in total). Among them, we selected the solution *C* such that *silh(C)* is maxima, which contained 18 clusters (*k* = 30, *r* = 0.7) and was computed on the top 20 PCs obtained from 399 features (mean.var.plot, d = 1.5; see “Generation of the clustering solution”). A tSNE transformation was computed, with perplexity = 10, on the same input used to compute *C*.

### Cluster labelling

Clusters identify was assigned by manual curation using the ImmGen MyGeneSet tool (http://rstats.immgen.org/MyGeneSet/). Clusters identified as thymic T cells, endothelial cells and epithelial cells were removed from downstream analyses, resulting in 14 differentiated immune cell clusters. 7395 cells were retained for WT and 7493 for Lamin A/C CKO.

### Differential expression analysis

Differential expression analysis was performed between each cluster and its complementary, and between the WT and *Lmna*^fl/fl^Vav1-Cre cell subsets, *cWT* and *cKO*, for each cluster *c* on solution *C,* using MAST method (Finak et al., 2015). Genes detected in ≥ 10% cells in either one of the two conditions with |log2FC|> 0.5 and adjusted p-value < 0.05 (Bonferroni correction) were defined as differentially expressed (DEGs).

For bootstrap validation, 10 cells subsets containing 50% of the cells were randomly generated for both cWT and cKO for each immune cell cluster *c* containing at least 50 cells in both cWT and cKO (12 out of the 14 immune cell clusters identified).. A DEG (|log2(FC)| > 0.5, adjusted p-value < 0.05, detected in ≥ 10% of the cells in either a subset or its complementary) found in at least one comparison between a subset and its complementary (within either *cWT* or *cKO*) was included in a *null set*. 100 cross-comparisons were performed between each pair of subsets generated from *cWT* and *cKO* and genes identified as DEGs are extracted (|log2(FC)| > 0.5, adjusted p-value < 0.05, detected in ≥ 50% of the cells in either one subset). A DEG detected in the full *cWT* versus *cKO* comparison was labelled as *validated* if it did not belong to the null set, was detected in ≥ 50% of the cells in either *cWT* or *cKO*, and was found among the top 25% significant DEGs in ≥ 90% of the cross-comparisons.

### Single-cell RNA-Seq of irradiated murine AM

Lungs were resected and single cell suspensions were prepared by enzymatic dissociation before loading into the Chromium controller (10x Genomics). Single-cell RNA-Seq of irradiated murine AM were prepared using single cell 3’RNA-Seq V2 reagent kit. cDNA quality control was assessed by capillary electrophoresis (Bioanalyzer, Agilent) before libraries preparation and sequencing on HiSeq 2500 (Illumina). Initial data processing was performed using the Cell Ranger pipeline (v2.1 10x Genomics). Count matrices were processed with Seurat package v2.5. Briefly, gene counts matrices were filtered (nGene < 2500; percentage of mitochondrial genes < 0.07), normalized and aligned using the canonical correlation analysis method in Seurat. After dimensionality reduction, clusters were annotated based on known cell type markers and DEG analysis performed on selected AM cluster. DEGs were identified in AM 5 months post-17Gy irradiation vs non-irradiated controls as genes that satisfy | log2(FC )*| >* 0.5 with an adjusted p-value *<* 0.05 and are detected in at least 10% of the cells in either one of the two conditions. We obtained 141 up- and 216 down-regulated genes for 5 months irradiation compared to control.

### Public single-cell RNA-Seq datasets of murine AM

#### Tabula Muris Senis (TMS)

The 10x AM data were obtained from the of the Tabula Muris Senis dataset (The Tabula Muris Consortium et al., 2020) (“alveolar macrophage” from cell_ontology_class_reannotated (https://s3.console.aws.amazon.com/s3/buckets/czb-tabula-muris-senis/)) for each age (1, 3, 18, 21, 30 months). We clustered the AM (k = 20, r = 0.1) on the top 10 PC computed on all genes and across all cells in the gene-cell matrix and computed the average expression of AM markers (see above).

#### Aging atlas

The cell-type-resolved differential gene expression testing between age groups was obtained (Angelidis et al., 2019) and the DEGs between AM in aged (24 months) versus young mice (3 months) were extracted (see paragraph “Differential expression analysis”).

#### Lung connectome

The gene expression profile of human, mouse, rat, and pig (two samples per species) across lung cells was obtained (Raredon et al., 2019) (accession: GSE133747). Based on the available annotation, immune cells were extracted and the average LMNA log-normalized expression (see paragraph “Processing of the gene-cell expression matrix”) across samples was computed.

DEGs were converted to the corresponding official gene symbols (using limma v3.38.3 (Smyth, 2005)) before being possibly compared across datasets.

### Definition of the TMS aging signature in mouse AM

Differential expression analysis was computed on each age pair and the DEGs were extracted as in paragraph “Differential expression analysis”. The DEGs for each old (18, 21, 30 months) versus young (1, 3 months) comparison are extracted. Up-(down-)regulated DEGs in one old-young comparison that are also found either down-(up-)regulated in another old-young comparison, or up-(down-)regulated in the 3 months versus 1month comparison, are filtered out. The remainder DEGs were validated using a similar boostrap procedure as the one described in paragraph “Comparison of WT versus *Lmna*^fl/fl^Vav1- Cre in each cell population”, allowing to define a list of validated up- and down-regulated genes for each of the six old-young comparison. The TMS aging signature was defined as the union of the six validated DEG lists.

### Functional annotation

For a given pair of conditions, functional annotation was performed separately on the up-regulated and down-regulated DEGs (see paragraph “Differential expression analysis”). Each DEG was first converted to the corresponding official gene symbol (using limma) and then to the Entrez ID. GO-BP enrichment analysis is computed using all mouse genes having a GO annotation as background (with clusterProfiler v3.10.0 (Yu et al., 2012) and org.Mm.eg.db v3.7.0). Only GO terms containing ≥ 10 and ≤ 500 genes are considered. A cut-off of 0.1 on the adjusted p-value (Benjamini-Hochbergh) is set to define the significantly enriched GO terms.

Redundant terms (Wang similarity measure > 0.7) are removed by keeping the most significant representative.

### Gene set enrichment analysis

Consistency between a defined gene set (A) and a phenotype (B) was quantified using GSEA (Subramanian et al., 2005). A custom gene signature was defined as the set of up-regulated DEG between case and control in A. The phenotype was defined as the average log2FC between case and control in B and used to rank the genes. Only the genes detected in at least 10% of the cells in either case or control in the phenotype were considered for GSEA.

## Supplementary Tables

Table S1 Differentially expressed genes for cluster assignment.

Table S2 Expression of Lmna in the Lung Connectome dataset.

Table S3 Differentially expressed genes identified in clusters.

Table S4 Differentially expressed genes in the alveolar macrophage cluster of irradiated mice

Table S5 Differentially expressed genes in the alveolar macrophage cluster of Tabula Muris Senis.

**Table S6.** Sex of mice used in the study.

| <u>Figure 1</u> |  |
| --- | --- |
| A | 5 M WT<br>6 M Lamin A/C CKO<br>10 F WT<br>7 F Lamin A/C CKO<br>*sex not noted for 4 WT and 3 CKO data points |
| B & C | 2 F WT<br>2 F Lamin A/C CKO |
| D | Data shown in Figure 1A<br>+<br>4 F WT<br>4 M Lamin A/C CKO |
| E | Sex not noted |
| <u>Figure 2</u> |  |
| A | 2 F WT<br>2 F Lamin A/C CKO |
| B | 7 M WT<br>8 WT Lamin A/C CKO<br>5 F WT<br>5 F Lamin A/C CKO |
| C | 3 M WT<br>4 M Lamin A/C CKO<br>2 F WT<br>2 F Lamin A/C CKO |
| D | 7 M WT<br>8 M Lamin A/C CKO<br>2 F WT |
|  | 2 F Lamin A/C CKO<br>2 M cGAS KO<br>4 M Lamin A/C cGAS DKO<br>2 F cGAS KO<br>3 F Lamin A/C cGAS DKO<br>5 M WT<br>5 M Lamin A/C p53 DKO<br>2 F WT<br>1 F Lamin A/C p53 DKO |
| <u>Figure 3</u> |  |
| A & B | 1 F |
| C & D | representative images and data from 4 experiments in which BAL from 2 WT mice was pooled<br>exp 1 - 2 M<br>exp 2 - 2 M<br>exp 3 - 1 M, 1F<br>exp 4 - 2M |
| E | images represenative of 4 experiments in which BAL from 2 mice of each genotype was pooled<br>exp 1 - 2 M WT + 2 M Lamin A/C CKO<br>exp 2 - 2 M WT + 2 M Lamin A/C CKO<br>exp 3 - 1 M WT + 1 F WT + 2 F Lamin A/C CKO<br>exp 4 - 2 M WT + 2 M Lamin A/C CKO |
| <u>Figure 4</u> |  |
| A | 2 F WT<br>2 F Lamin A/C CKO |
| B | 10 M WT<br>13 M Lamin A/C CKO<br>7 F WT<br>6 F Lamin A/C CKO |
| C | 2 F WT<br>2 F Lamin A/C CKO<br>2 F WT Control<br>2 F WT Irradiated |
| D | 7 M WT<br>8 M Lamin A/C CKO<br>2 F WT<br>2 F Lamin A/C CKO<br>2 M cGAS KO<br>4 M Lamin A/C cGAS DKO<br>2 F cGAS KO<br>3 F Lamin A/C cGAS DKO<br>5 M WT<br>5 M Lamin A/C p53 DKO |
|  | 2 F WT<br>1 F Lamin A/C p53 DKO |
| <u>Figure 5</u> |  |
| A | not noted |
| B | 3 F WT<br>2 M Lamin A/C CKO<br>1 F Lamin A/C CKO |
| <u>Figure 6</u> |  |
| A & B | 12 F WT<br>12 F Lamin A/C CKO |
| <u>Figure 7</u> |  |
| A & B | data from Tabula muris senis |
| C | 8F |
| D & E | 2 F WT<br>2 F Lamin A/C CKO<br>and data from Tabula muris senis and Lung Aging Atlas |
| <u>Figure S1</u> |  |
|  | Sex not noted |
| <u>Figure S2</u> |  |
| A & B | 5 M WT<br>6 M CKO<br>8 F WT<br>5 F CKO<br>*sex not noted for 4 WT and 3 CKO data points |
| C | sex not noted |
| <u>Figure S3</u> |  |
| <u>A-C</u> | 2 F WT<br>2 F Lamin A/C CKO |
| <u>Figure S4</u> |  |
| A & B | 2 F WT<br>2 F Lamin A/C CKO |
| C | representative example of data shown in Figure 2D and 4D |
| <u>Figure S5</u> |  |
| A | 1F |
| B | representative example of data shown in Figure 3C, D and E |
| C | representative images and data from 1 experiment in which BAL was pooled from 2 mice of each genotype<br>2 M WT |
|  | 2 M Lamin A/C CKO |
| D | representative images and data from 3 experiments in which BAL from 2 mice of each genotype was pooled<br>exp 1 - 1 M WT + 1 F WT + 2 M Lamin A/C CKO<br>exp 2 - 1 M WT + 1 F WT + 2 M Lamin A/C CKO<br>exp 3 - 1 M WT + 1 F WT + 1 M Lamin A/C CKO + 1 F Lamin A/C CKO |
| <u>Figure S6</u> |  |
| A | 1 M WT<br>3 M Lamin A/C CKO<br>4 F WT<br>5 F Lamin A/C CKO |
| B | 6 M WT<br>6 M Lamin A/C CKO<br>4 F WT<br>5 F Lamin A/C CKO |
| C | 3 M WT<br>5 M Lamin A/C CKO<br>5 F WT<br>2 F Lamin A/C CKO |
| D | 2 F WT Control<br>2 F WT Irradiated |
| E | 3 F WT<br>2 M Lamin A/C CKO<br>1 F Lamin A/C CKO |
| <u>Figure S7</u> |  |
| A | 12 F WT<br>12 F Lamin A/C CKO |
| B-D | 3 F WT<br>3 M WT<br>3 F Lamin A/C CKO<br>3 M Lamin A/C CKO |
| <u>Figure S8</u> |  |
| A & C | 8 F |
| B | 2 F WT<br>2 F Lamin A/C CKO<br>and data from Tabula muris senis |

**Table S7.**
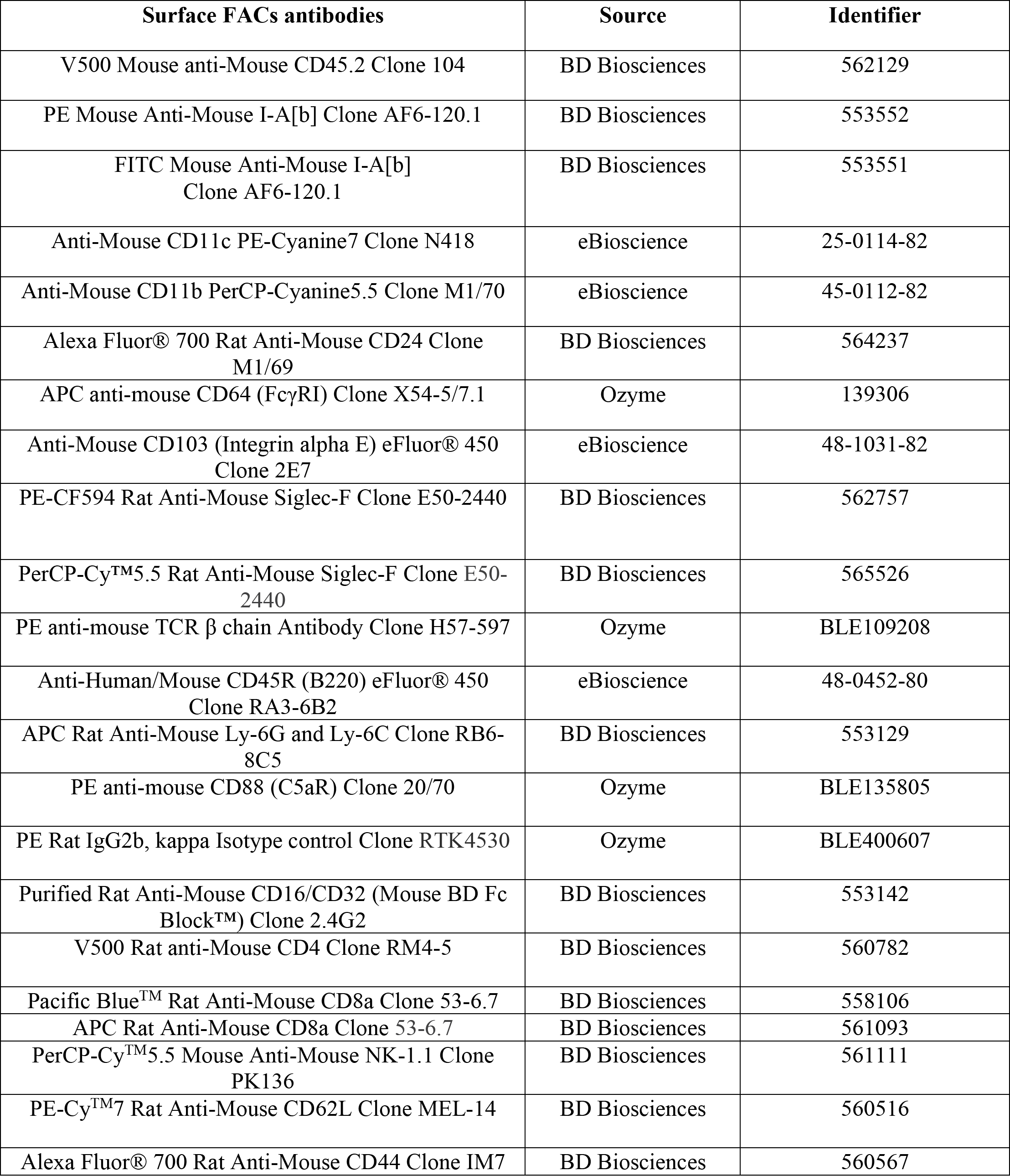
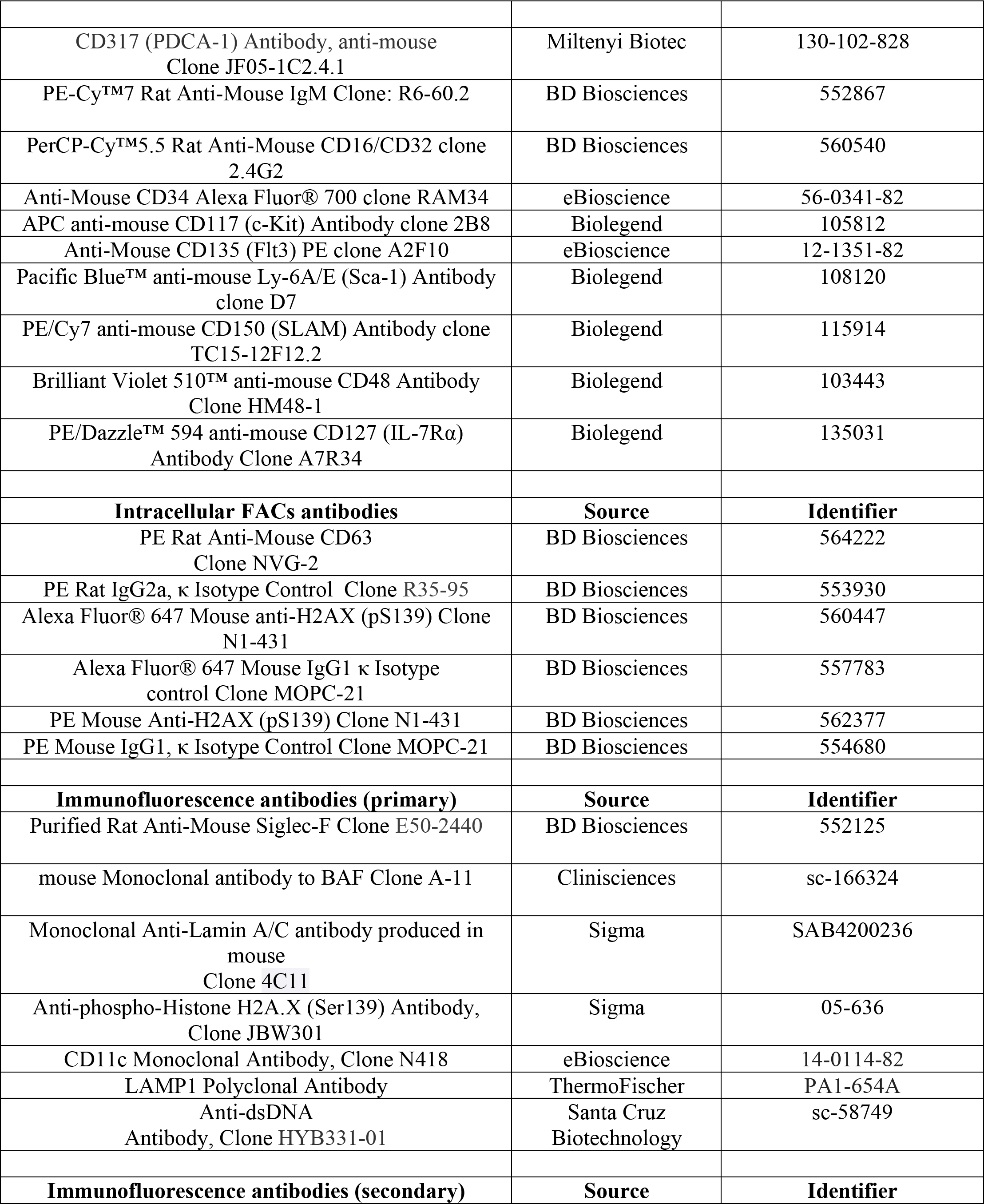
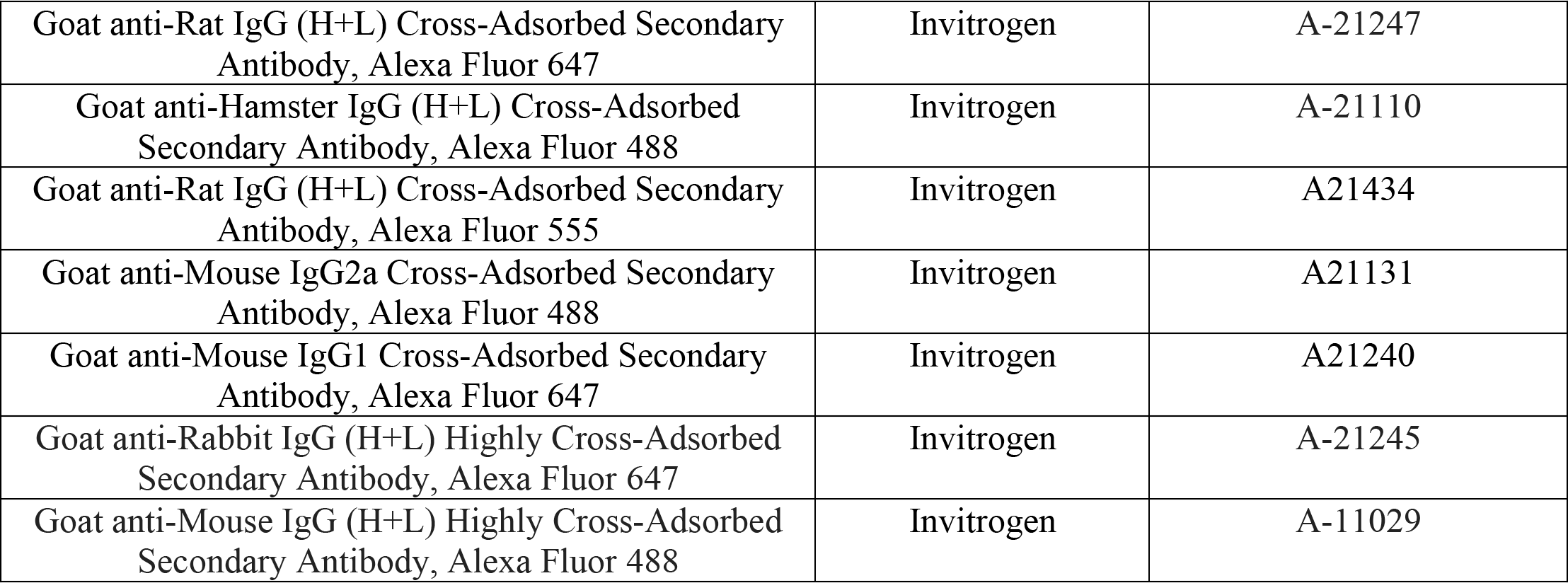
Antibodies used in the study.

**Supplementary figure 1.**
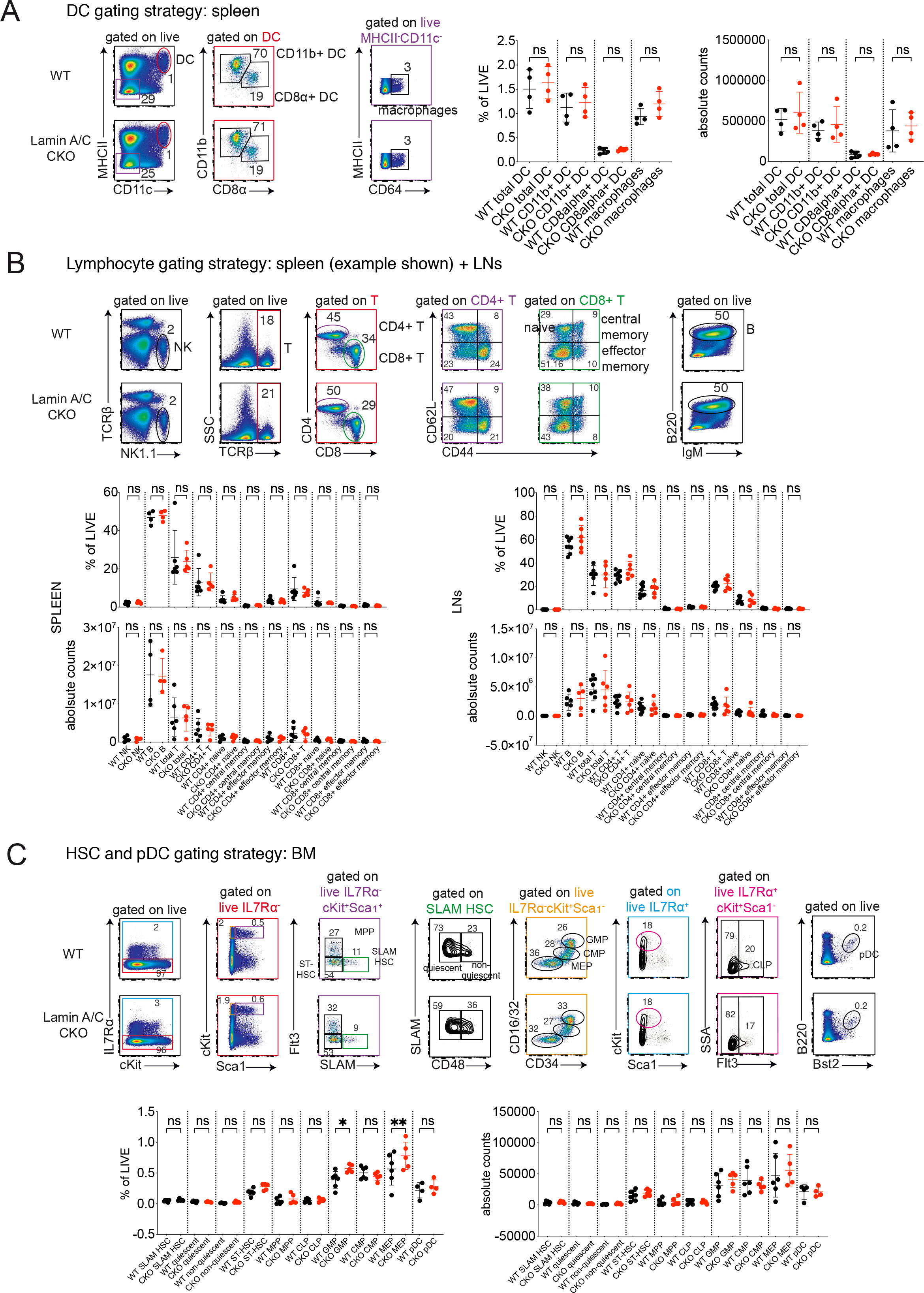
Analysis of immune cells in Lamin A/C CKO mice. **(A)** Flow cytometric analysis of WT vs Lamin A/C CKO macrophage and dendritic cell (DC) populations in spleen, with gating strategies for each population represented by colored gating. Left, representative example of the gating strategy. Right, percentage of each population of live spleen cells and absolute counts (n = 4 mice, combined from 2 independent experiments, bar indicates mean ± SD, one-way ANOVA with Šidák test). **(B)** Flow cytometric analysis of WT vs Lamin A/C CKO Nature killer, T and B cell populations in spleen and lymph nodes, with gating strategies for each population represented by colored gating. Top, representative example of the gating strategy. Bottom, percentage of each population of live spleen or lymph node cells and absolute counts (n = 5-8 mice, combined from 4 independent experiments, bar indicates mean ± SD, one-way ANOVA with Šidák test). **(C)** Flow cytometric analysis of WT vs Lamin A/C CKO to analyze hematopoietic stem cell (HSC) and plasmacytoid dendritic cell populations in bone marrow, with gating strategies for each population represented by colored gating. Top, representative example of the gating strategy. Bottom, percentage of each population of live bone marrow cells cells and the absolute counts (n = 4-6 mice, combined from 3 independent experiments, bar indicates mean ± SD, one-way ANOVA with Šidák test). Acronyms used: Short term HSC (ST-HSC), Multipotent hematopoietic progenitor (MPP), common myeloid progenitor (CMP), common lymphoid progenitor (CLP), granulocyte macrophage progenitor (GMP), megakaryocyte-erythroid progenitor (MEP), plasmacytoid dendritic cell (pDC).

**Supplementary figure 2.**
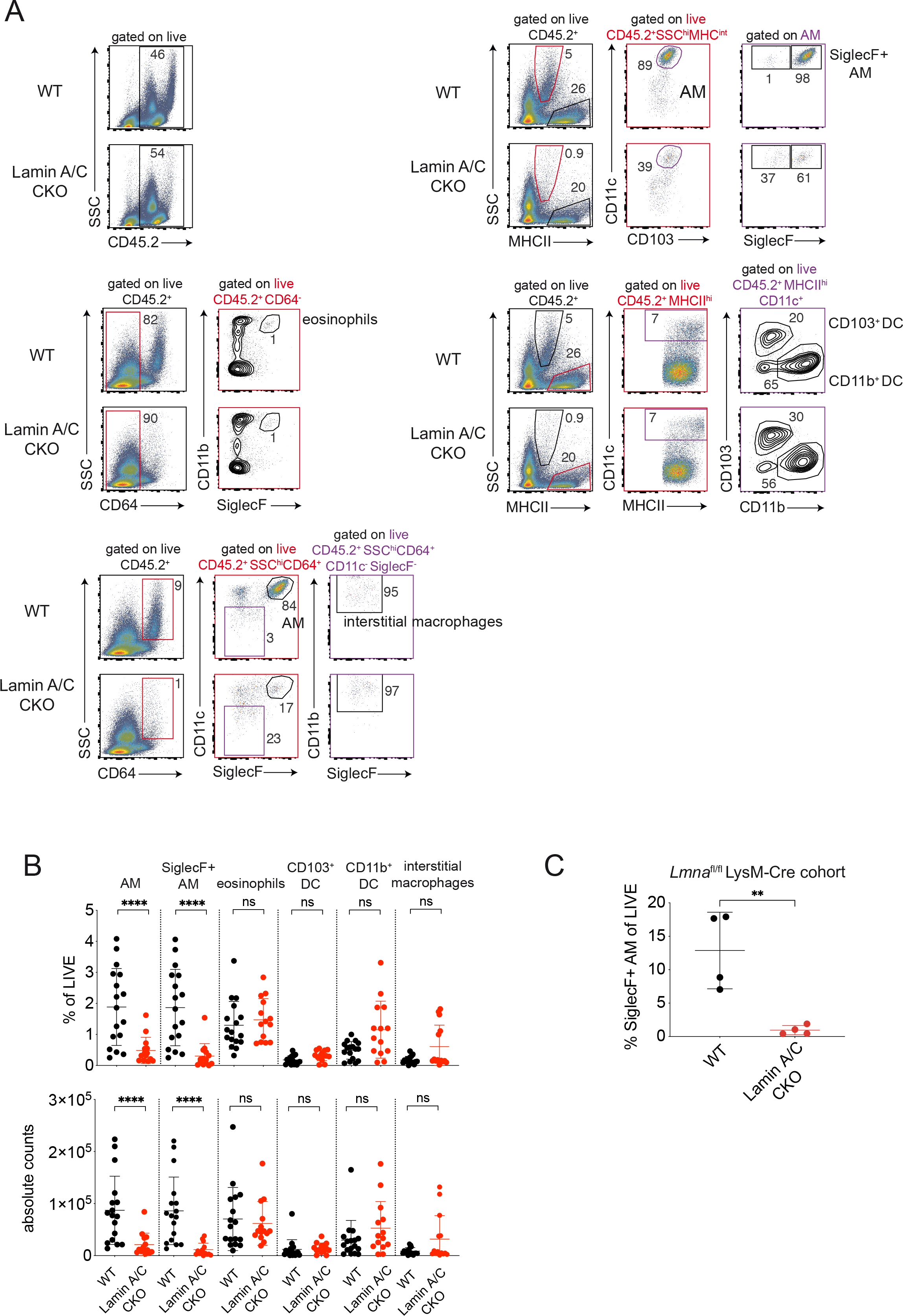
Analysis of immune cells in lungs of Lamin A/C CKO mice. **(A)** Representative flow cytometric analysis of WT vs Lamin A/C CKO lung immune populations including AM, eosinophils, interstitial macrophages, CD103+ DC and CD11b+ DC with gating strategies for each population represented by colored gating. **(B)** Percentages and absolute counts of AM, eosinophils, interstitial macrophages, CD103+ DC and CD11b+ DC of live WT vs Lamin A/C CKO lung cells (n = 14-17 mice, combined from 9 independent experiments, bar indicates mean ± SD, one-way ANOVA with Šidák test). **(C)** Percentage of AM (CD45.2^+^SSC^hi^MHC^int^SiglecF^+^) following flow cytometric analysis of WT (*Lmna*^fl/fl^ LysM-Cre^-/-^) vs *Lmna*^fl/fl^ LysM-Cre^+/-^ lung (n = 4 mice combined from 2 independent experiments, bar indicates mean ± SD, unpaired t test).

**Supplementary figure 3.**
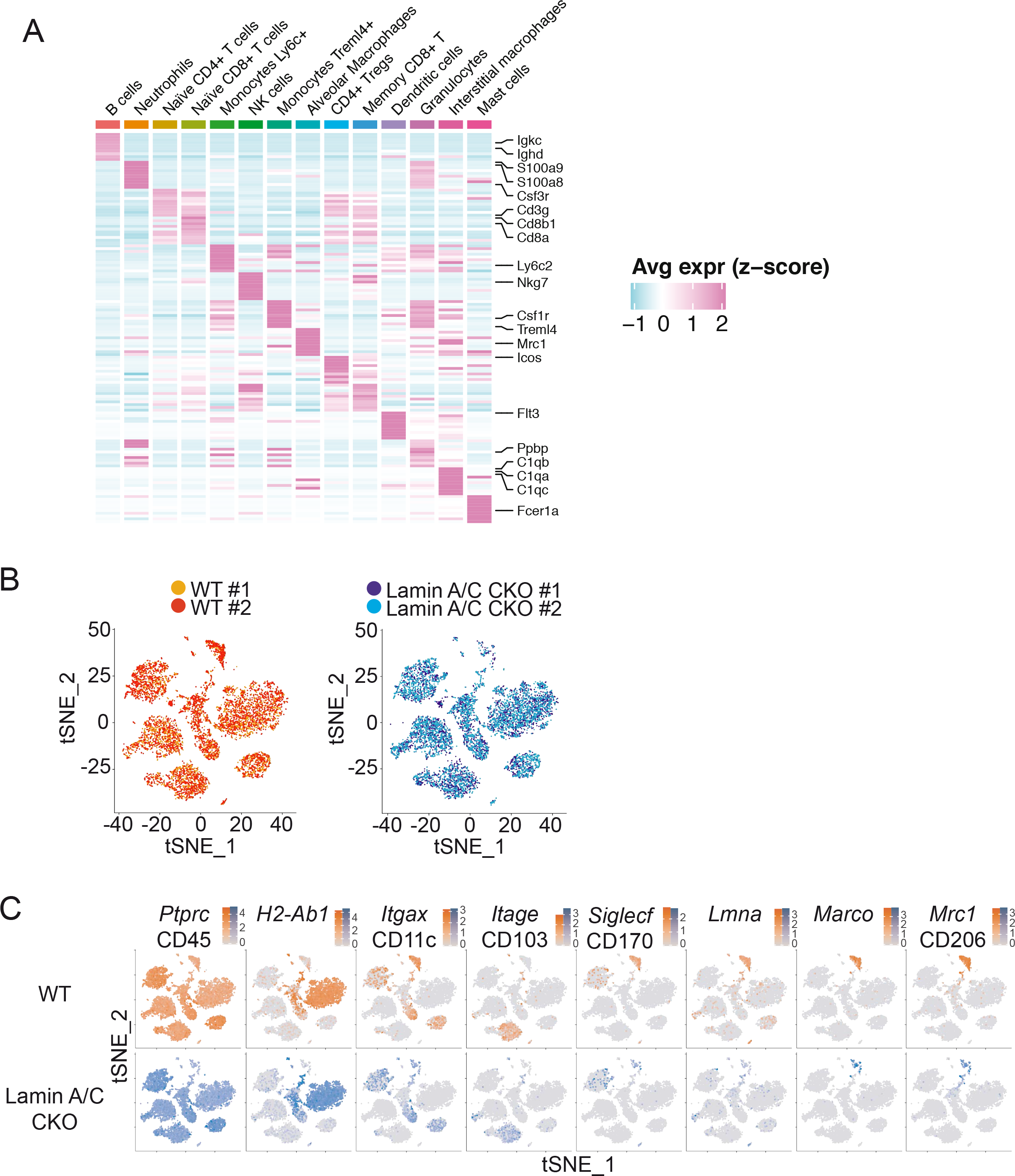
scRNAseq analysis of lung immune cells in WT and Lamin A/C CKO mice. **(A)** Heatmap showing the expression of cluster-specific gene markers. Differential expression analysis is done for each cluster against all the other cells (including the contaminants). Among the top 50 significant DEGs, the 10 DEGs with the top fold change are shown. Common DEGs between clusters are reported only once. Entries are the z-scores of the normalized expression level, clipped to fit the interval. **(B)** tSNE representation showing WT (left) and Lamin A/C CKO (right) cells only, and colored by replicate. In WT, replicate 1 contains 4075 cells and replicate 2 contains 3980. In Lamin A/C CKO, replicate 1 contains 4000 cells and replicate 2 contains 3677. **(C)** tSNE representation showing WT cells (top) and Lamin A/C CKO cells (bottom). Cells are colored with a gradient representing the normalized expression level of immune and alveolar macrophage markers.

**Supplementary figure 4.**
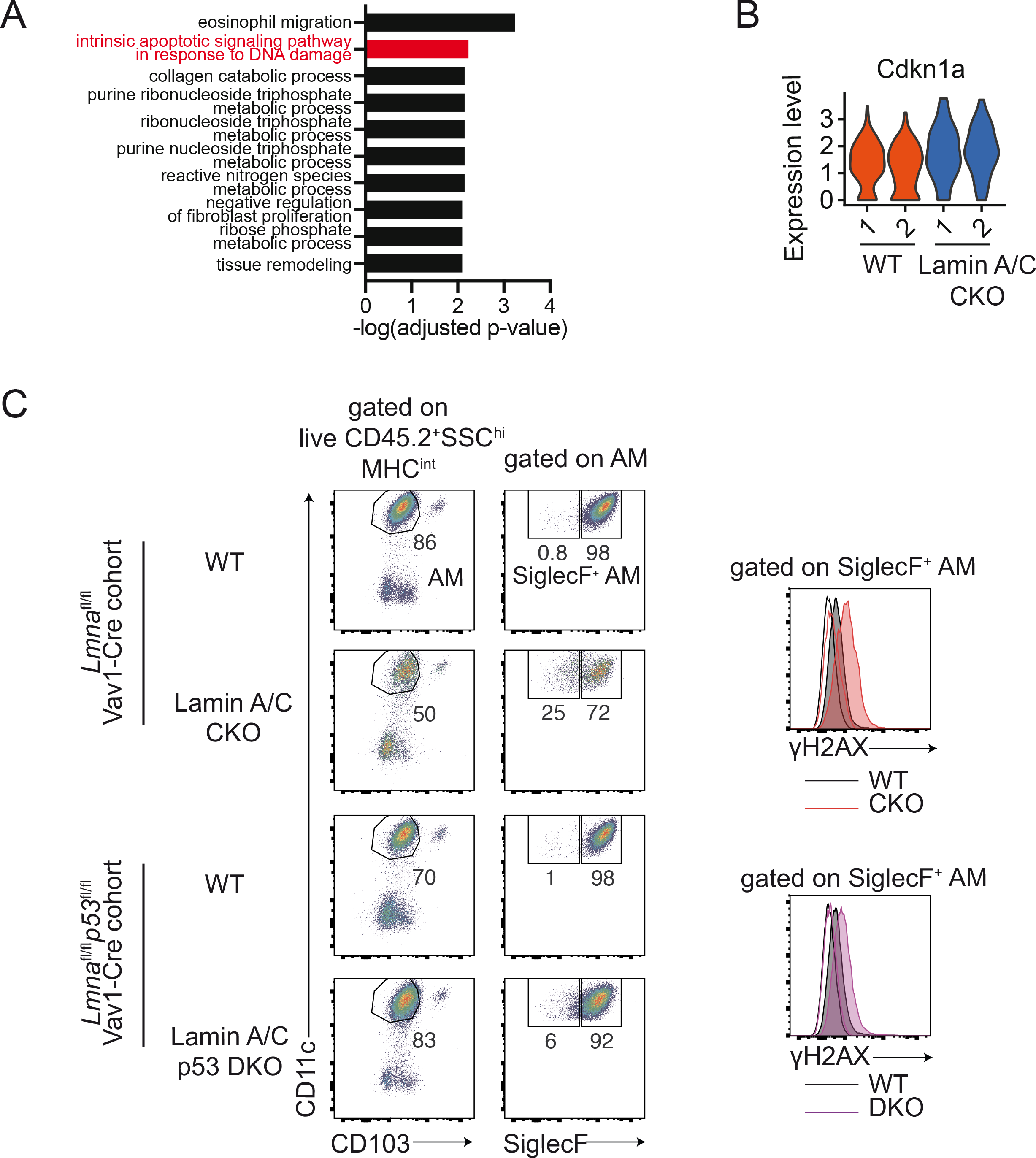
DNA damage and related markers in alveolar macrophages of Lamin A/C CKO mice. **(A)** Gene ontology enrichment analysis of DEGs upregulated in Lamin A/C CKO, specific to the AM cluster. Significant terms (adjusted p-value < 0.01) from the biological process (BP) ontology are shown. **(B)** Violin plot showing the normalized expression of *Cdkn1a* in the AM cluster, with values for individual WT and Lamin A/C CKO replicates shown. **(C)** Representative intracellular flow cytometric analysis of mice from WT (*Lmna*^fl/fl^ Vav1-Cre^-/-^) vs Lamin A/C CKO (*Lmna*^fl/fl^ Vav1-Cre^+/-^) and and WT (*Lmna*^fl/fl^ *p53*^fl/fl^Vav1-Cre^-/-^) vs Lamin A/C p53 DKO (*Lmna*^fl/fl^ *p53*^fl/fl^Vav1-Cre^+/-^) to identify AM (CD45.2^+^SSC^hi^MHC^int^SiglecF^+^) and assess γH2AX levels.

**Supplementary figure 5.**
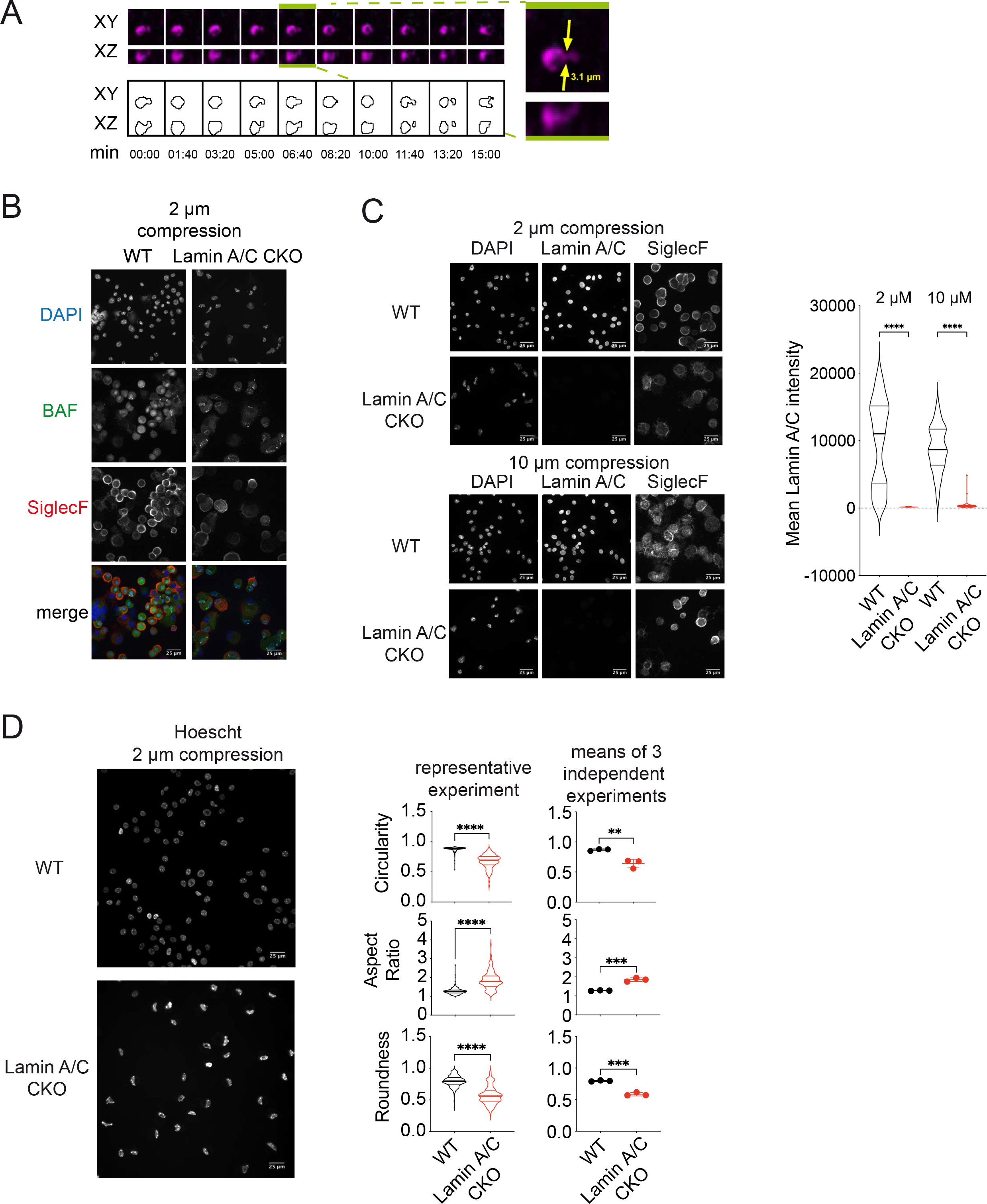
Live imaging of alveolar macrophages in lungs. **(A)** AM from WT mouse lung (female) undergoing constricted migration *in vivo*. Top left, XY and XZ planes. Bottom left, contour representation. Right, measurement of the cell width at the most acute point of squeezing. **(B)** BAL from Lamin A/C CKO mice confined at a height of 2 μm for 1.5 hours and subsequently stained for DAPI, BAF and SiglecF (representative of n = 4 independent experiments, each time BAL was pooled from 2 mice aged 8-26 weeks). **(C)** BAL from WT and Lamin A/C CKO mice confined at 2 µm and 10μm for 1.5 hours and subsequently stained for DAPI, Lamin A/C and SiglecF. Left, representative field. Right, quantification of mean intensity of the Lamin A/C intensity (violin plot with lines indicating the median and quartile values, representative of 2 independent experiments, one-way ANOVA with Tukey test, each time BAL was pooled from 2 mice of each genotype). **(D)** Quantification of circularity, aspect ratio and roundness of nuclei from WT and Lamin A/C CKO BAL. BAL was stained with Hoescht, confined at 2 μm and imaged immediately. Left, representative field. Middle, values of individual cells in one experiment (violin plot with lines indicating the median and quartile values, Kolmogorov-Smirnov test). Right, means of 3 independent experiments (unpaired t-test). For each experiment BAL was pooled from 2 mice of each genotype aged 10-27 weeks. In the experiment shown, the cells were also stained for CD11c and SiglecF prior to confinement to quantify nuclear shape specifically in AM. Bars indicate mean ± SD.

**Supplementary figure 6.**
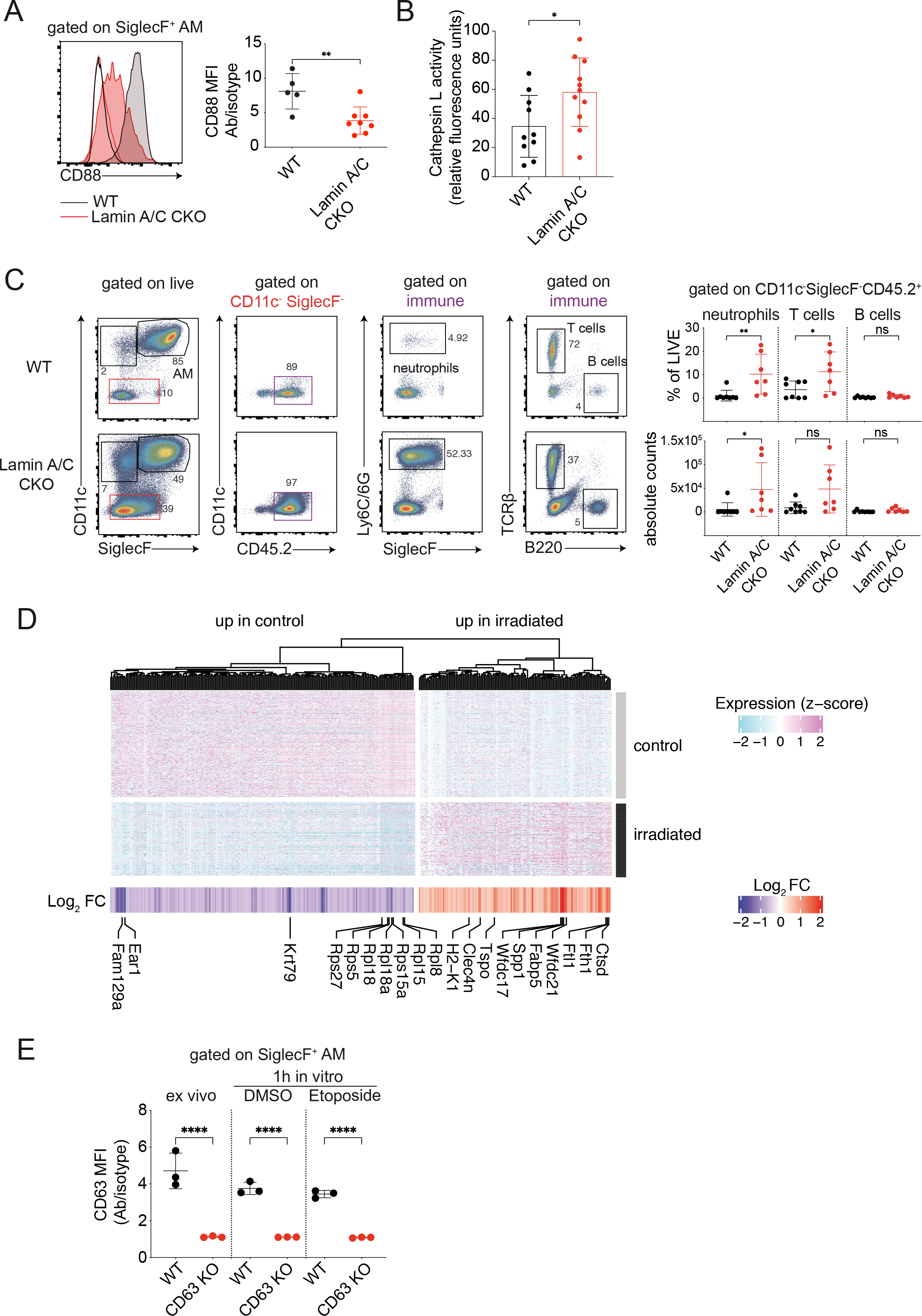
Analysis of markers in Lamin A/C CKO mice, scRNAseq of irradiated mice and CD63 expression in knock-out mice. **(A)** Left, intracellular flow cytometric analysis of WT vs Lamin A/C CKO lung for CD88 levels within AM (CD45.2^+^SSC^hi^MHC^int^SiglecF^+^). Right, mean fluorescence intensity (MFI) of CD88 signal normalized to isotype control (n = 5-8 mice, combined from 3 independent experiments, mice aged 28-39 weeks, bar indicates mean ± SD, unpaired t test). **(B)** Cathepsin L activity in BAL extracted from WT vs Lamin A/C CKO mice (n = 10-11 mice from 8 independent experiments, bar indicates mean ± SD, aged 13-34 weeks, unpaired t-test). **(C)** Flow cytometric analysis of BAL obtained from WT vs Lamin A/C CKO mice to analyze neutrophils, T and B cells with gating strategies for each population represented by colored gating. Left, representative example. Right, percentage of each population of live BAL cells and absolute counts of each population in extracted BAL (n = 7-8 mice, combined from 4 independent experiments, mice aged 24-34 weeks, bars indicate mean ± SD, one-way ANOVA with Šidák test). **(D)** Heatmap showing single cell gene expression (z-score) of DEGs in AM 5 months post-17Gy irradiation and non-irradiated controls. Cells are grouped by condition. DEGs are divided among up- and down-regulated in case vs control, clustered with a ward.D2 method computed on euclidean distances and shown as dendrogram. DEGs are annotated with log2 fold change (FC) values. **(E)** Intracellular flow cytometric analysis for CD63 levels within AM (CD45.2^+^SSC^hi^MHC^int^SiglecF^+^) of WT vs CD63 KO lung single cell suspension, treated as indicated. MFI of CD63 normalized to isotype control (n = 3 mice in one experiment, mice were 19 weeks of age, bars indicate mean ± SD, one-way ANOVA with Šidák test).

**Supplementary figure 7.**
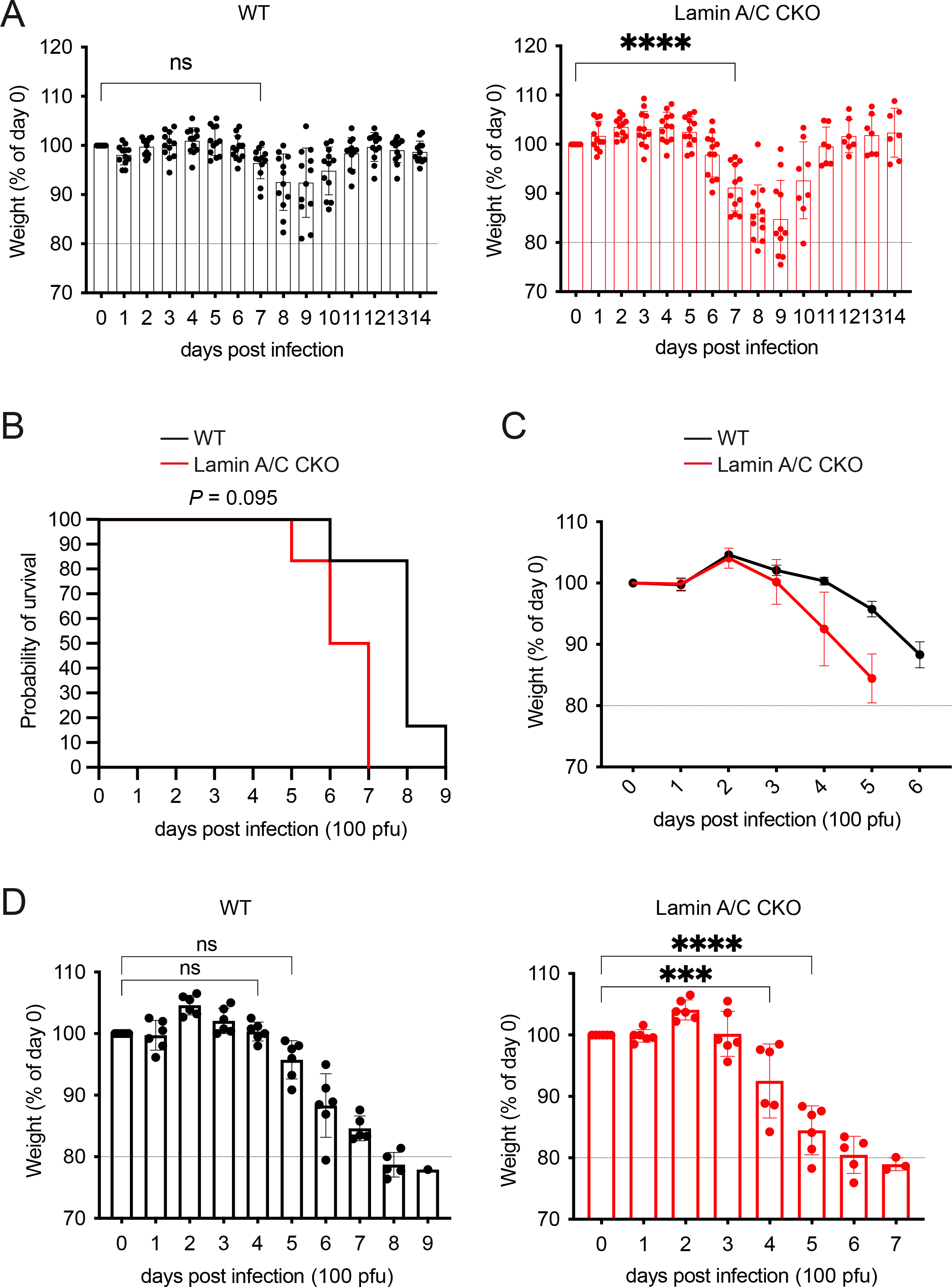
Influenza infections in Lamin A/C CKO mice. **(A)** Percentage of day 0 weight observed each day post infection with 50 pfu influenza A virus via the intranasal route in WT and Lamin A/C CKO mice (n = female 12 mice per genotype, combined from 2 independent experiments, two-way ANOVA with Tukey test, bars indicate mean ± SD). **(B)** Survival of WT vs Lamin A/C CKO mice following infected with 100 pfu of influenza A virus PR8 via the intranasal route (n = 6 mice per genotype in one experiment, Log-rank Mantel-Cox test). **(C)** Percentage of day 0 weight observed each day post infection described in (B). Curves of average ± SEM weights are shown and continued until a first death occurs in the group. **(D)** Percentage of day 0 weight observed each day post infection described in (B) (two-way ANOVA with Tukey test, bars indicate mean ± SD).

**Supplementary figure 8.**
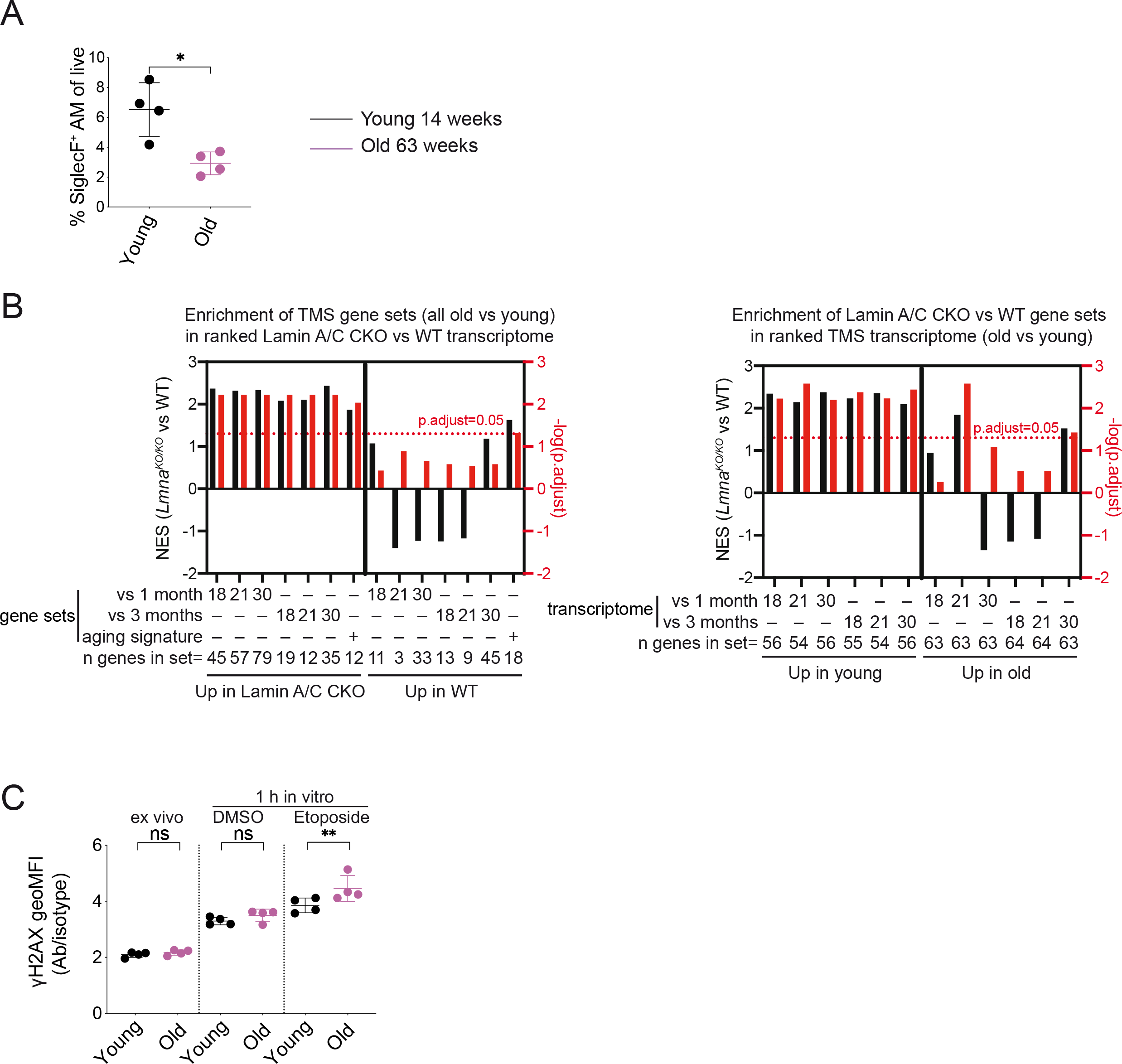
Hallmarks of aging in alveolar macrophages. **(A)** Percentage AM (CD45.2^+^SSC^hi^MHC^int^SiglecF^+^) of live lung cells in Young (14 weeks) vs Old (63 weeks) mice (n = 4 mice combined from 2 independent experiments, bars indicate mean ± SD, unpaired t test). **(B)** Normalized enrichment scores (NES) and adjusted p-values following gene set enrichment analysis of the indicated upregulated or downregulated genes. Left, TMS gene sets enrichment in Lamin A/C CKO vs WT transcriptome. Geneset obtained after individual pairwise time-point comparisons and overall aging signature were tested. Right, Lamin A/C CKO gene sets enrichment in TMS transcriptome, ranked by the indicated time-point comparisons. n genes in sets indicates the number of genes in given gene set detected in the tested transcriptome. **(C)** Young (14 weeks) vs Old (63 weeks) lung single cell suspension was directly analyzed or incubated with DMSO or 50 μM etoposide for 1 hour. Intracellular flow cytometric analysis for γH2AX levels within AM (CD45.2^+^SSC^hi^MHC^int^SiglecF^+^) was then performed. The geoMFI of γH2AX signal is shown normalized to isotype control (n = 4 mice from 2 independent experiments, bars indicate mean ± SD, one-way ANOVA with Šidák test).

Movie 1 Live imaging of a wild-type mouse lung.

The lung was imaged after administration of Hoechst and an anti-SiglecF antibody. Left is a broad field of a lung region, right features individual AM (SiglecF^+^) demonstrating constricted migration (white boxes).

Movie 2 A single AM undergoing constricted migration *in vivo*.

Left, contour representation. Right, XY and XZ planes.

